# Derivation of functional early gestation decidual natural killer cell subtypes from induced pluripotent stem cells

**DOI:** 10.1101/2025.11.03.685424

**Authors:** Virginia Chu Cheung, Jennifer Jaimez, Carly DaCosta, Harneet Arora, Christine Caron, Jaroslav Slamecka, Manuel Fierro, Morgan Meads, Kathleen Fisch, Robert E. Morey, Luisjesus S. Cruz, Devika Pant, Dan S. Kaufman, Mariko Horii, Jack D. Bui, Mana M. Parast

## Abstract

Abnormal decidual natural killer cell (dNK) function is linked to pregnancy complications occurring in both early and late gestation, including recurrent pregnancy loss, preeclampsia, and preterm birth. Exploration of dNK heterogeneity as it relates to function is an active area of research; however, most of this work has focused on early gestation. Using flow cytometric and transcriptomic single-cell definitions of dNK subtypes, we characterized dNK heterogeneity in term dNK within both chorioamniotic membranes and basal plate. We also applied aptamer-based secretome profiling to first trimester and term dNK, and found dNK-specific proteins – VEGF and PLGF – to be reduced at term. We further determined that, compared to first trimester dNK, term dNK have reduced cytoxicity against target cells. Finally, we applied this knowledge to establish a protocol for differentiation of induced pluripotent stem cells (iPSC) into functional dNK. We found that treatment with TGFβ enriched for dNK2 subtype, while inducing dNK markers, CD9 and CD103. We evaluated function using cytokine and degranulation assays, aptamer-based secretome profiling, and cytotoxicity assays. We found that iPSC-dNK are functionally most similar to primary dNK. Further, TGFβ iPSC-dNK had reduced GM-CSF in response to PMA/I and increased secretion of VEGF and other first trimester-specific proteins – supportive of a shift towards an early gestation, dNK2-dominant, phenotype. We conclude that changes in dNK function across gestation reflect shifts in dNK subtypes that can be reproducibly derived from iPSC, providing a new method for modeling dNK and laying the foundation for cell-based therapeutics for reproductive disease.

**Significance Statement:** Alterations in maternal decidual natural killer cells (dNK) are associated with pregnancy complications – from recurrent pregnancy loss to preeclampsia and preterm birth. We found dNK from different regions of the term placenta to be distinct from peripheral blood NK and early gestation dNK, based on gene and surface marker expression, subtype composition, secretome, and cytotoxicity. We report a novel, reproducible protocol to generate dNK resembling the most abundant dNK subtype in early gestation from induced pluripotent stem cells. Our study lays the foundation for *in vitro* modeling of the maternal-fetal interface and therapeutic development for reproductive disease.

## Introduction

A successful pregnancy requires maternal tolerance of the semi-allogeneic fetal-derived placenta.^1^ Prior to fertilization and embryo implantation, the uterine lining undergoes decidualization, a morphologic and functional transformation, leading to a tissue poised for both tolerance of fetal-derived antigens, and recognition and clearance of potential pathogens.^1,2^ This process involves infiltration with a vast array of maternal leukocytes, including natural killer (NK) cells.^3^ Decidual NK cells (dNK) are a type of tissue resident NK cell (trNK) found in the uterine lining. dNK cells function to support implantation and promote angiogenesis by crosstalk with maternal decidualized endometrial stromal fibroblasts (dESF) and fetal-derived extravillous trophoblasts (EVT) in the placenta.^4,5^ As gestation progresses, the maternal decidua contributes to the basal plate (BP, also the decidua basalis) and chorioamniotic membranes (CAM, also the decidua parietalis). dNK at the BP regulate EVT invasion and establish maternal blood flow to the feto-placental unit.^5^ Meanwhile, CAM functions as a protective barrier surrounding the fetus throughout much of gestation. Abnormal dNK function has been associated with multiple pregnancy complications including recurrent spontaneous pregnancy loss, preeclampsia, as well as inflammation-induced spontaneous preterm birth.^6–9^

dNK have cell surface marker expression and function distinct from peripheral blood NK cells (PB-NK). dNK in nonpregnant and first trimester decidua are CD56^bright^CD16^-^, and co-express CD9, unlike CD56^dim^CD16^+^CD9^-^ PB-NK.^10,11^ In the NK-92 cell line, acquisition of CD9 results in increased production of proangiogenic IL-8, reduced production of proinflammatory TNFα and IFNγ, and reduced cytotoxicity against target cells.^12^ However, dNK at the maternal-fetal interface are not the same throughout gestation. Inhibitory Killer Cell Immunoglobulin-Like Receptors KIR2DL1 and KIR2DL2/3 are reduced in both BP- and CAM-dNK cells at term compared to first trimester dNK.^13,14^ At baseline, dNK and PB-NK express cytolytic molecules (PFN, GZMB, GNLY); however, PB-NK have higher PFN and GZMB than dNK and first trimester dNK (but not term dNK) have higher GNLY and 9kD GNLY.^13^ Unstimulated, first trimester dNK, but not PB-NK, secrete proangiogenic cytokines and growth factors (IL-6, VEGF, PLGF).^15,16^ These data are supportive of a role for dNK in vascular remodeling in early gestation, however, the function of unstimulated term dNK have never been evaluated in this capacity.

Primary dNK have reduced cytotoxicity relative to PB-NK; however, this function is context-dependent and regulated by the decidual niche. *In vitro* first trimester dNK demonstrate reduced lysis of target cells in comparison to PB-NK, but first trimester dNK with high IL-15 will lyse target cells more than PB-NK.^17^ In addition, first trimester dNK have been shown to lyse K562 and primary trophoblast target cells similar to PB-NK, but this is diminished in the presence of decidual macrophages or TGFβ.^18^ While term dNK degranulate in response to phorbol- 12-myristate-13-acetate and ionomycin (PMA/I) stimulation and K562 at levels similar to PB-NK, term dNK have different expression of cytolytic proteins relative to both PB-NK and first trimester dNK.^13^ Thus, it remains unclear if term dNK have the capacity to kill target cells.

Nevertheless, it is known that PB-NK, first trimester dNK, and term dNK cells have distinct activation responses. PB-NK induce higher levels of CD107a and cytokines (TNFα, IFNγ, GM-CSF), compared to first trimester dNK, in response to PMA/I stimulation.^13,15^ Given the same stimulation, term dNK (BP and CAM) respond with high CD107a, similar to PB-NK, yet have a modest pro-inflammatory cytokine response (TNFα, IFNγ), similar to first trimester dNK. Although certain activation receptors (*CD69*, *NKp80*, *NKp44*) are expressed at the RNA level in CAM-dNK, but not in BP-dNK, functional and protein expression differences have not yet been identified.^13,19–21^ dNK function and receptor expression are distinct between first trimester and term (including between BP and CAM), however, the basis for this functional change remains unclear.

Recent single-cell analyses have pointed to first trimester dNK’s transcriptional and functional heterogeneity, identifying 4 subtypes, dNK1, dNK2, dNK3 and dNKp.^22–24^ dNK1 express CD39 (*ENTPD1*), an inhibitory receptor expressed by trNK found in tonsil, liver, and first trimester decidua, CD39 knocked out mice have increased NK cell numbers and NK effector functions.^11,25^ dNK2 and dNK3 express CD18 (*ITGB2*), an integrin that enhances NK cell cytotoxicity.^26^ dNK3 also express CD103 (*ITGAE*), a tissue residency marker for lung NK cells, as well as T cells in multiple tissue types.^27,28^ Expression of CD103 by dNK is associated with higher cytokine (GM-CSF, XCL1) and CD107a expression, after PMA/I stimulation.^23^ Following stimulation, dNK3 have the highest GM-CSF expression followed by dNK2, then dNK1 with low levels similar to PB-NK.^23,24^ Importantly, dNK subtypes differentially express receptors and ligands (*CCL5*, *LILRB1*, *CSF1*) used for interaction with placental EVT and other cell types– suggesting distinct roles in placentation.^22^ However, it is not known how changes in these subtypes relate to functional changes throughout gestation at different locations within the maternal-placental interface.

Due to lack of access to decidual tissues in an ongoing pregnancy, studies rely on model systems. Mouse placentation is distinct from human at structural, cellular, and molecular levels.^29^ Importantly, human and mouse placental cells at the maternal-placental interface do not have the same expression of key receptor/ligand pairs involved in dNK crosstalk.^30^ Similarly, PB-NK and term dNK significantly differ from their first trimester counterparts, both in their immunophenotype and functional capacity.^3,13, 31–33^ While attempts have been made to generate dNK from PB-NK, primary NK cells are difficult to reliably expand in culture – limiting their capacity for mechanistic studies.^34,35^ This necessitates novel models to study human dNK and their integral interactions with maternal and placental cells.

Over the past two decades, generation of patient-specific induced pluripotent stem cells (iPSC) from somatic cells has revolutionized the modeling of numerous human diseases.^36^ We and others have shown that preeclampsia (PE)-associated cellular abnormalities can be modeled using iPSC. This includes abnormal differentiation and function of trophoblast, derived from iPSC, obtained by reprogramming cells from PE-affected placentas,^37–39^ as well as abnormal function of endothelial cells, derived from iPSC, obtained by reprogramming peripheral blood mononuclear cells (PBMCs) from patients with PE.^40^ Currently, there are protocols to routinely generate mature, cytotoxic CD56^+^CD16^+^ NK cells from iPSC. These iPSC-NK transcriptionally resemble umbilical cord and PB-NK and kill tumor cells *in vitro* and *in vivo.*^41–43^ Several clinical trials have now demonstrated that engineered iPSC-NK are safe and effective in treatment of refractory malignancies.^36,44^ However, no protocol exists to generate dNK from iPSC.

Studies in mice and human suggest one mechanism of dNK emergence to be through PB-NK infiltration and exposure to the decidual niche,^45,46^ where the oxygen tension throughout gestation is 2-8% O_2_.^47^ In fact, IL-15 and exposure to low oxygen tension induce a dNK-like phenotype in PB-NK.^34,35^ Trophoblast-derived SDF-1 (*CXCL12*) stimulate PB-NK migration into decidua, and PB-NK coculture with EVT has been shown to promote conversion to a dNK-like phenotype.^45,48^ Furthermore, treatment of PB-NK with TGFβ1, a growth factor secreted by maternal dESF and placental EVT, in combination with hypoxia, promote a dNK-like phenotype, including induction of CD9, reduced cytotoxicity, and increased VEGF secretion.^27,35^ In this study, we characterize dNK heterogeneity and function at first trimester and different regions of the term placenta, and apply this improved understanding of dNK gene expression, surface protein and cytokine expression, protein secretion, and cytotoxicity, to establish a differentiation protocol to derive functional dNK from iPSC.

## Materials and Methods

For detailed protocols, see **Supplementary Materials and Methods.**

### Patient recruitment and tissue collection

Human decidual and blood tissue samples were collected under UCSD IRB-approved protocols (IRB #181917 and 172111); all patients gave informed consent.

### hPSC

Human pluripotent stem cell (hPSC) experiments were performed under protocol #171648, approved by the UCSD IRB and Embryonic Stem Cell Research Oversight Committee. Human embryonic stem cell (hESC) WA09/H9 and 3 iPSC lines (established as a part of the biorepository of the Center for Perinatal Discovery) were used for differentiation.

## Results

### Defining primary dNK subtypes by scRNA-seq and flow cytometry across gestation

We set out to identify first trimester dNK subtypes (dNK1, dNK2, dNK3, and dNKp) in term decidua.^22,23^ We reanalyzed previously-published scRNA-seq datasets from blood and first trimester decidua (Vento-Tormo, 2018) as well as term placental and decidual tissues (Pique-Regi, 2019).^22,49^ We utilized all cells in the Vento-Tormo dataset, and verified NK cells by expression of NK-associated genes (**Supplementary Figure 1A, B**). We extracted all the NK cells (peripheral blood, PB-NK; and decidua, dNK) (**Figure 1A**), to use as a reference. We compared gene expression between these reference NK cells, in the absence of non-NK cell types, to identify new NK cell subtype-specific marker genes for dNK1, dNK2, dNK3, dNKp, PB-NK CD16^+^, and PB-NK CD16^-^ (**Figure 1A**, **Supplementary Figure 1C**, **Table 1, Supplementary File 1)**.

**Figure 1.**
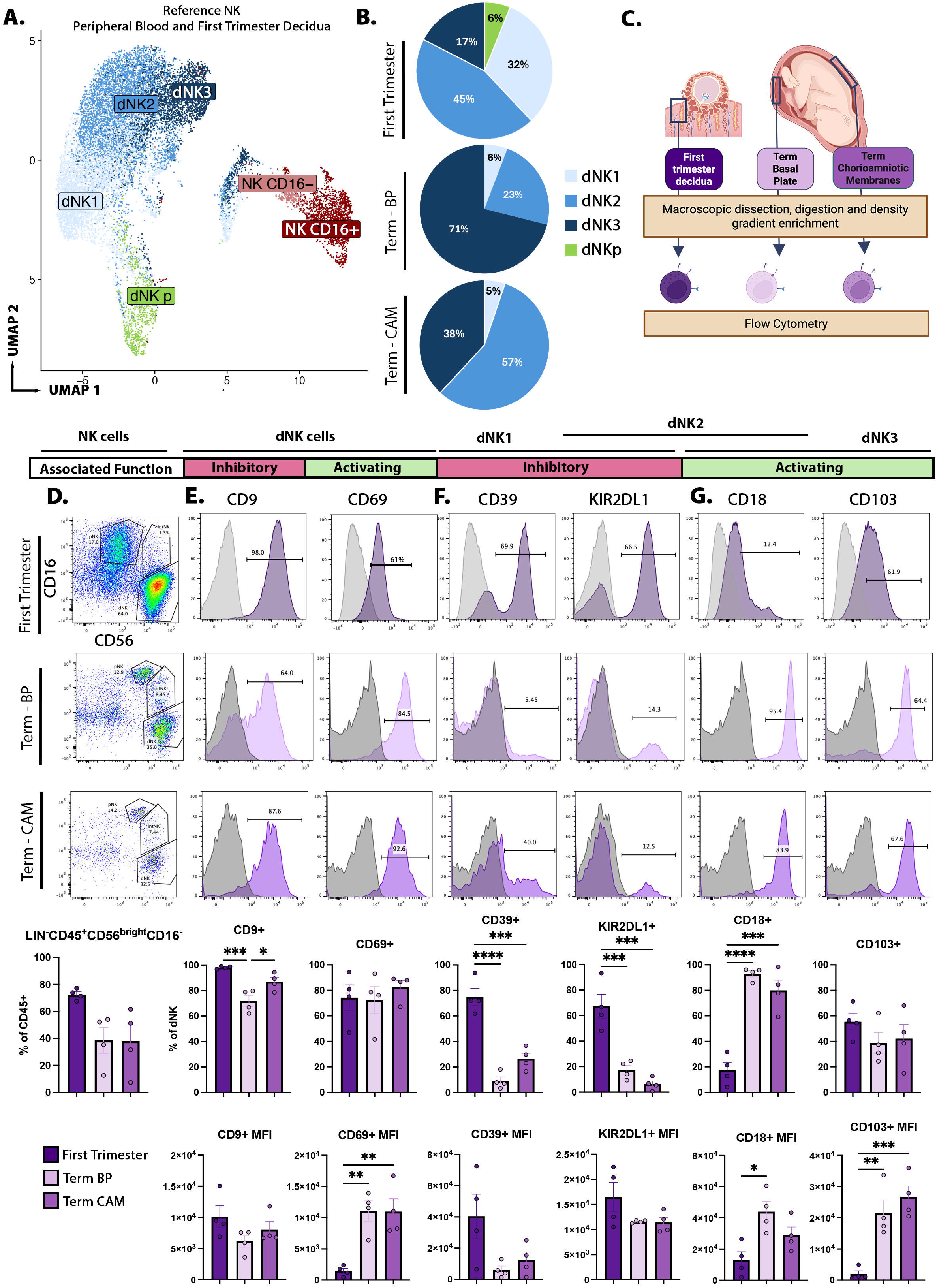
Term dNK cells show enrichment in activated dNK subtypes. **A)** UMAP showing NK cell reference from Vento-Tormo, 2018^22^ containing dNK and PB-NK subtypes. **B)** Pie charts showing dNK subtype composition in decidua from first trimester, term basal plate (BP), and term chorioamniotic membrane (CAM), as identified by reference mapping of term scRNA-seq data from Pique-Regi et al., 2019^49^ to **(A)** (data from Pique-Regi reanalyzed with permission). **C)** Schematic of primary cell samples used for flow cytometric analysis. **D-G)** Representative FACS plots and quantification of NK cell markers (CD56 and CD16), in first trimester decidua, term BP, and term CAM after gating on Live Lin^-^CD45^+^ **(D)**, as well as dNK markers (CD9 and CD69) **(E)**, inhibitory receptors (CD39 and KIR2DL1) **(F)**, and activating receptors (CD18 and CD103) **(G)** after gating on Live Lin^-^CD45^+^CD56^bright^CD16^-^ cells. For **D** through **G**, data are represented as mean +/- standard error, n = 8 individual patients (4 first trimester donors and 4 term placenta donors; BP and CAM were dissected and processed separately from the same term placentas). Statistical analysis was performed using ordinary one-way ANOVA with multiple comparisons using GraphPad Prism; *p<0.05, **p<0.01, ***p<0.001, ****p<0.0001. Alt text: UMAP showing NK cell heterogeneity. Pie charts showing dNK subtype composition across gestation. Histograms and bar graph quantification of cell surface marker expression on first trimester, Term BP, and Term CAM dNK cells.

**Table 1.** Top 25 marker genes distinguishing dNK cell subtypes (dNK1, dNK2, dNK3, dNKp) and PB-NK cell subtypes (PB-NK CD16^+^, PB-NK CD16^-^)

| **dNK1** | **dNK2** | **dNK3** | **dNKp** | **PB-NK**<br>**CD16-** | **PB-NK**<br>**CD16+** |
| --- | --- | --- | --- | --- | --- |
| *B4GALNT1* | *CDHR1* | *ITM2C* | *PBK* | *IL7R* | *FGFBP2* |
| *CYP26A1* | *CD2* | *CXCR4* | *CENPA* | *SELL* | *S1PR5* |
| *ENTPD1* | *ZNF683* | *TAGAP* | *CDCA3* | *IL18R1* | *AKR1C3* |
| *SPINK2* | *COX6A2* | *CD160* | *PLK1* | *IGFBP4* | *SPON2* |
| *TPTEP1* | *TNFRSF4* | *LINC01871* | *DLGAP5* | *RIPOR2* | *PRSS23* |
| *C15orf48* | *KRT81* | *RGCC* | *KIF23* | *LTB* | *FCGR3A* |
| *EPAS1* | *CTSA* | *PDE4B* | *UBE2C* | *PLAC8* | *ADGRG1* |
| *CDKN1A* | *IFI44L* | *DUSP4* | *PIMREG* | *TXNIP* | *PLAC8* |
| *CSF1* | *C1orf162* | *ARL4A* | *CCNA2* | *LINC00861* | *S100A8* |
| *UBASH3B* | *IGFBP1* | *RGS2* | *HMMR* | *RASGRP2* | *LINC00861* |
| *UBE2F* | *ITGAX* | *DNAJB1* | *SPC25* | *GZMK* | *TBX21* |
| *DAPK2* | *BCO2* | *CEMIP2* | *TOP2A* | *IGKC* | *RIPOR2* |
| *ACY3* | *SNRNP25* | *RASGEF1B* | *CEP55* | *TCF7* | *LYAR* |
| *SYNGR1* | *KIR2DL4* | *ZNF331* | *RRM2* | *MCUB* | *GNG2* |
| *ID3* | *IGFBP2* | *IFRD1* | *AURKB* | *KLF2* | *CD160* |
| *PLPP1* | *MATK* | *NFE2L2* | *CKAP2L* | *GRK6* | *TXNIP* |
| *MDM2* | *TSPO* | *ZEB2* | *CDCA5* | *KLRF1* | *CLEC2D* |
| *STAT3* | *AGTRAP* | *CCNH* | *CDK1* | *SNHG25* | *S100A9* |
| *CD59* | *TESC* | *PIK3R1* | *CCNB2* | *LINC01871* | *C1orf21* |
| *SLC12A8* | *SLC44A1* | *PTPN22* | *SPC24* | *ITGAM* | *ICAM2* |
| *AFAP1L2* | *LASP1* | *PMAIP1* | *KIF2C* | *S100A9* | *TTC38* |
| *PIK3R6* | *AHNAK* | *ALOX5AP* | *GTSE1* | *TRGC2* | *FAM107B* |
| *CAMK1* | *SEPTIN11* | *TIPARP* | *CDCA8* | *ATM* | *C12orf75* |
| *BCL2L11* | *GZMH* | *SERTAD1* | *NUSAP1* | *ARHGAP15* | *LBH* |
| *SRGAP3* | *CHCHD10* | *ITGAE* | *BIRC5* | *VPS51* | *PPP2R5C* |

In the Pique-Regi dataset, three compartments in the term placenta (BP; CAM; and placental villus, PV) were annotated using sample-associated meta data (**Supplementary Figure 1D,E**). We verified NK cells by expression of NK-associated genes (**Supplementary Figure 1F**). We extracted all NK cells from the Pique-Regi dataset and mapped them to our reference NK cells.^22^ All dNK subtypes found in first trimester could be identified at term, with dNK2 making up the majority of the dNK in first trimester decidua (45%) and term CAM (57%) while dNK3 make up the majority in term BP (71%) (**Figure 1B**). Meanwhile, dNK1 made up only 6% of BP-dNK and 5% of CAM-dNK, while <0.1% of all term dNK mapped to dNKp (**Figure 1B**). We applied our new NK cell subtype- specific markers (**Supplementary Figure 1C**) to the term NK dataset and found an abundance of cells expressing dNK3 marker, *CCL5*, with smaller populations of cells expressing the dNK2 marker, *ZNF683* (also known as HOBIT), and dNK1 marker, *SPINK2*, and only rare cells expressing the dNKp marker *MKI67* (**Supplementary Figure 1C,G,H, Table 1**). We found there were more dNK2 marker, *ZNF683^+^* NK cells in CAM and more dNK3 marker*, CCL5^+^* NK cells in BP (**Supplementary Figure 1I**). This agrees with the reference mapping results, which identified more dNK2 in CAM and more dNK3 in BP (**Figure 1B**). The Pique-Regi dataset contains cells from placentas delivered at term, with and without labor, as well as those delivered preterm with labor.^49^ Across all three labor groups, dNK3 were increased in both compartments relative to first trimester, with dNK3 dominant in BP, and dNK2 dominant in CAM (**Supplementary Figure 1J.**).

We next characterized dNK in early first trimester decidua and term placenta, the latter including decidua from both BP and CAM, by flow cytometry (**Figure 1C**). We confirmed that dNK are CD45^+^CD56^bright^CD16^-^ and comprise 40-70% of CD45^+^ cells in first trimester and term decidual tissues (**Figure 1D**). dNK-associated CD9 and trNK-associated CD69 were expressed by a large proportion of dNK throughout gestation (**Figure 1E**).^12,20^ CD9^+^ dNK was significantly reduced, 1.4-fold in term BP (71.9% mean) and 1.2-fold in CAM (87.0% mean), compared to first trimester. Quantification by Median Fluorescent Intensity (MFI) showed a significant increase in CD69 in term dNK in both compartments (**Figure 1E**).

We next evaluated dNK subtype-associated markers. dNK1 marker CD39 (*ENTPD1*) was expressed by an average of 74.9% of dNK in first trimester but was reduced over 8-fold (9.0% mean) in term BP and almost 3-fold (26.5% mean) in term CAM (**Figure 1F**). ^22,24^ Similarly, KIR2DL1, a marker of both dNK1 and dNK2, was expressed by an average of 67.2% of first trimester dNK, but only 17.5% and 6.4% of term BP and CAM, respectively (**Figure 1F**). ^22,23^ In contrast, CD18 (*ITGB2*), a marker of both dNK2 and dNK3 was ∼5-fold higher in term BP (93.0% mean) and term CAM (80.0% mean), compared to first trimester (17.6% mean) (**Figure 1G**). dNK3 marker CD103 (*ITGAE*) was expressed by an average of ∼55%, ∼39%, and 42% of the dNK in early gestation decidua, term BP, and term CAM, respectively (**Figure 1G**). ^22,23^ However, like CD18, CD103 MFI was upregulated at term, increased by ∼11-fold and 13.5-fold in term BP and CAM, respectively (**Figure 1G**). This is consistent with scRNA-seq results, which showed that dNK1 are more abundant in first trimester decidua, while dNK3 are relatively more abundant at term, in both BP and CAM. Flow cytometry of dNK2-associated KIR2DL1 and CD18 were incongruous with each other; however, scRNA-seq analyses using marker gene and unsupervised (reference mapping) analyses pointed to dNK2 as the most abundant subtype in first trimester and term CAM.

Term BP- and CAM-dNK have greater expression of activating receptors and degranulation response than first trimester dNK.^13^ When first trimester dNK is sub-stratified by subtype, dNK3 has the highest degranulation and cytokine expression (effector response), followed by dNK2, and then dNK1 upon PMA/I stimulation.^23,24^ In agreement with the literature, we found markers associated with decreased effector function (CD9, CD39 and KIR2DL1) to be higher in first trimester dNK, while activating receptors (CD69, CD18, and CD103) were increased at term, in both BP and CAM (**Figure 1E - G**). Interestingly, CD39, KIR2DL1, and CD18 were dramatically changed between first trimester and term. CD9, CD69, and CD103 were maintained by a more constant proportion of dNK, although levels of CD69 and CD103 were higher at term, compared to first trimester. Together, these data suggest that functional changes in dNK across gestation may be driven by a shift in dNK subtypes, with decreased dNK1 (low effector function, high levels of inhibitory molecules) and increased dNK3 (high effector function, high levels of activating molecules) as gestation progresses toward term. Additionally, we identify the proportion of dNK2 and CD9^+^ dNK to be similar between first trimester and term CAM.

### iPSC-NK acquire dNK markers in response to TGFβ treatment

We next set out to establish an *in vitro* model of dNK, by differentiation of pluripotent stem cells (both embryonic and induced PSC). We used a protocol, modified from Zhu et al., 2019,^50^ to differentiate the hESC line H9/WA09, and 3 different iPSC lines, generated and validated in our own lab (**Supplementary Figure 2A-E**), into iPSC-NK. At the start of this protocol (**Figure 2A**), hematopoietic stem/progenitor cells were generated from highly pure, SSEA4^+^ hPSC (**Figure 2B, C**), using a spin embryoid body (EBs) method. EBs containing CD34^+^ cells (**Figure 2B, D**) were plated for NK differentiation. After 28 days, round cells emerged in suspension. An average of 87.3% of these cells were CD45^+^, of which 78.2% were also CD56^+^ (**Figure 2B, E**). To enrich for NK, these cells were collected for expansion. Instead of coculture with artificial antigen presenting cells as previously published,^50^ we cultured the cells in a commercially available media (see **Methods**) supplemented with IL-2 and IL-15 for this expansion phase. After 2 weeks of expansion, an average of 87.8% of CD45^+^ cells were also CD56^+^, with 15.6% CD56^+^CD16^+^, and 72.2% CD56^+^CD16^-^ (**Figure 2B, F**).

**Figure 2.**
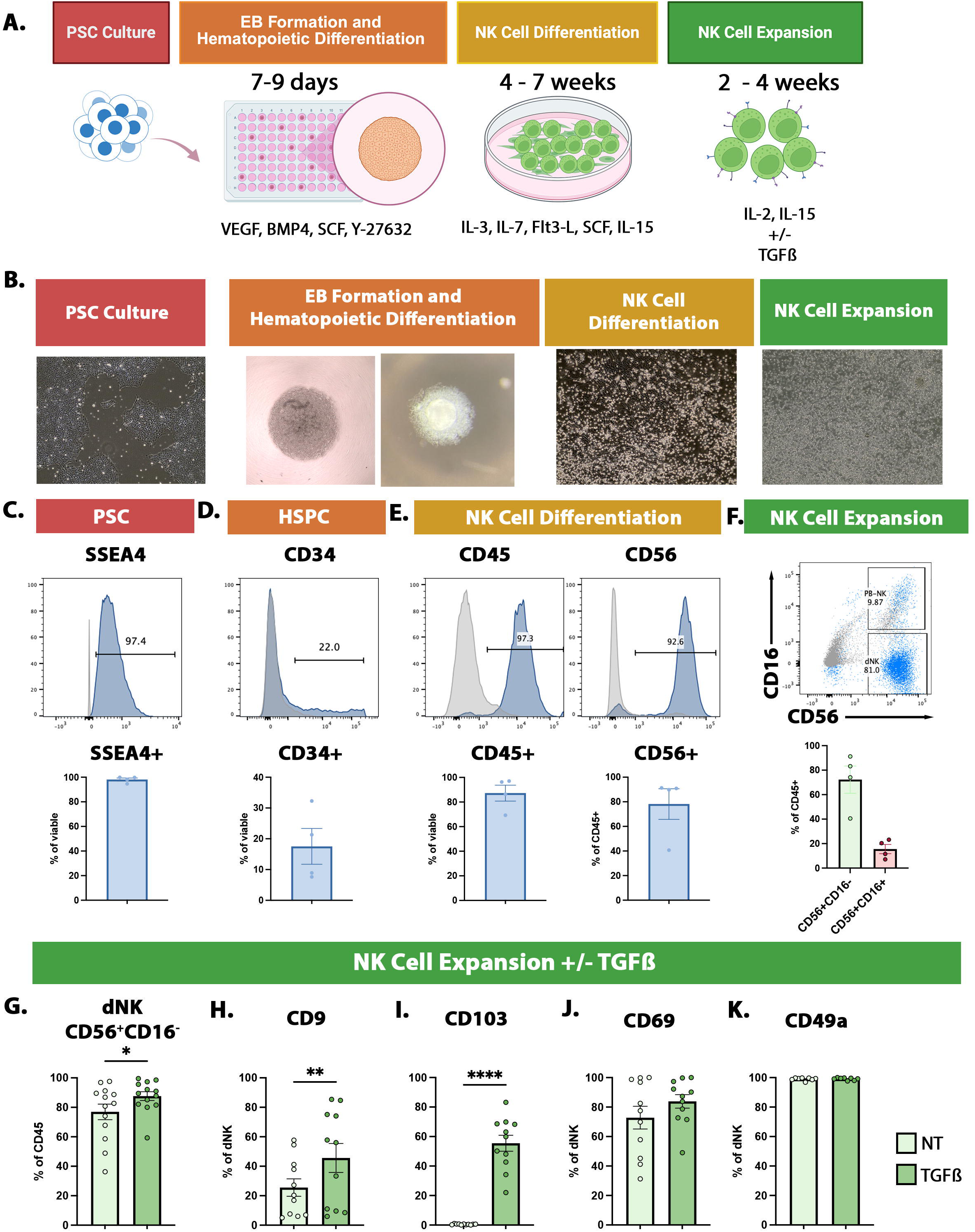
TGFβ supplementation increases dNK cell marker expression in iPSC-NK. **A)** Schematic of the entire differentiation protocol. **B)** Representative phase images of cells throughout the differentiation process. **C-E)** Representative FACS plot and quantification of expression over isotype control. Pluripotency marker SSEA4 staining in iPSC, before starting the differentiation **(C)**, hematopoietic stem progenitor cell (HSPC) marker CD34 staining after EB formation, before plating for NK cell differentiation **(D)**, and immune cell marker CD45 and NK cell marker CD56 staining after NK cell differentiation, before plating for expansion **(E)**. **F)** Representative FACS plot of CD56 and PB-NK marker CD16 co-staining after at least 2 weeks of expansion in IL-2 and IL-15. CD56^+^CD16^-^ and CD56^+^CD16^+^ populations are quantified. **G)** Quantification of CD45^+^CD56^+^CD16^-^ population in No Treatment (NT, no TGFβ) and TGFβ treatment conditions. **H-K)** Quantification of flow cytometric staining of dNK associated markers, CD9 **(H)**, CD103 **(I)**, CD69 **(J)**, and CD49a **(K)**, in iPSC-NK at the end of the expansion phase. For **C** through **K**, data are represented as mean +/- standard error. For **C** through **F**, n = 4 PSC lines; H9 hESC line and MB3140, MB3144, LD2809 iPSC lines. For **G** through **K,** n = 11 independent experiments using MB3140, MB3144, LD2809 iPSC lines, Statistical analysis was performed using Ordinary one-way ANOVA with multiple comparisons using GraphPad Prism; *p<0.05, **p<0.01, ***p<0.001, ****p<0.0001. Alt text: Schematic of iPSC differentiation protocol. Bright phase images of cells throughout differentiation. Histograms and bar graphs quantifying expression of markers during key phases of differentiation. Scatter plot and bar graphs showing dNK-like and PB-NK-like cells. Bar graphs showing increased dNK marker expression in iPSC-NK after TGFbeta treatment.

We hypothesized conditions mimicking the decidual niche would promote a dNK phenotype. We reclustered the NK cells from the Vento-Tormo dataset broadly into two groups: dNK (comprised of dNK1, dNK2, dNK3, and dNKp), or PB-NK (comprised of NK CD16^+^ and NK CD16^-^) (**Supplementary Figure 3A**) and compared the two to identify an enrichment in TGFβ signaling and hypoxia pathways in dNK (**Supplementary Figure 3B**). Additionally, TGFβ and CXCL12 have been shown to promote dNK-like phenotype when applied to PB-NK.^34,35,45^ We tested TGFβ and CXCL12 under 5% or 21% O_2_, during the expansion phase of our protocol. We found TGFβ, but not CXCL12 or 5% O2, slightly increased the proportion of CD56^+^CD16^-^, decreased CD18, and significantly increased dNK markers CD9 (2-fold) and CD103 (∼90-fold) (**Figure 2G, H, I, Supplementary Figure 3C, D**). TGFβ had no effect on CD69 and dNK-associated CD49a (**Figure 2J,K**).

Together, these data show that we can derive NK cells from hPSC, and that TGFβ pushes the cells further toward a dNK phenotype.

### iPSC-NK are transcriptionally similar to dNK, not PB-NK, and TGFβ induces the dNK2 phenotype

We further characterized our iPSC-NK, with and without TGFβ, using scRNA-seq (**Figure 3A**). We found that iPSC-NK expressed dNK marker genes (*NCAM1*, *ITGA1*, *CD69*) and no *FCGR3A* (PB-NK-associated CD16) (**Figure 3B**). We again used the full Vento-Tormo dataset as a reference, with NK cells subclustered broadly as dNK (comprised of dNK1, dNK2, dNK3, and dNKp), or PB-NK (comprised of NK CD16^+^ and NK CD16^-^) (**Supplementary Figure 3A**).^22^ We found 94% of iPSC-NK cells mapped to dNK (**Figure 3C,D**). iPSC-NK also expressed dNK-(*SPINK2*, *ZNF683*, *CCL5*), but not PB-NK- (*SELL*, *SPON2*) associated markers (**Supplementary Figure 4A**). These data indicate that the iPSC-NK, regardless of TGFβ treatment, transcriptionally resemble dNK, not PB-NK; henceforth, we will refer to our cells as iPSC-dNK, no TGFβ treatment as NT iPSC-dNK, and TGFβ treated cells as TGFβ iPSC-dNK.

**Figure 3.**
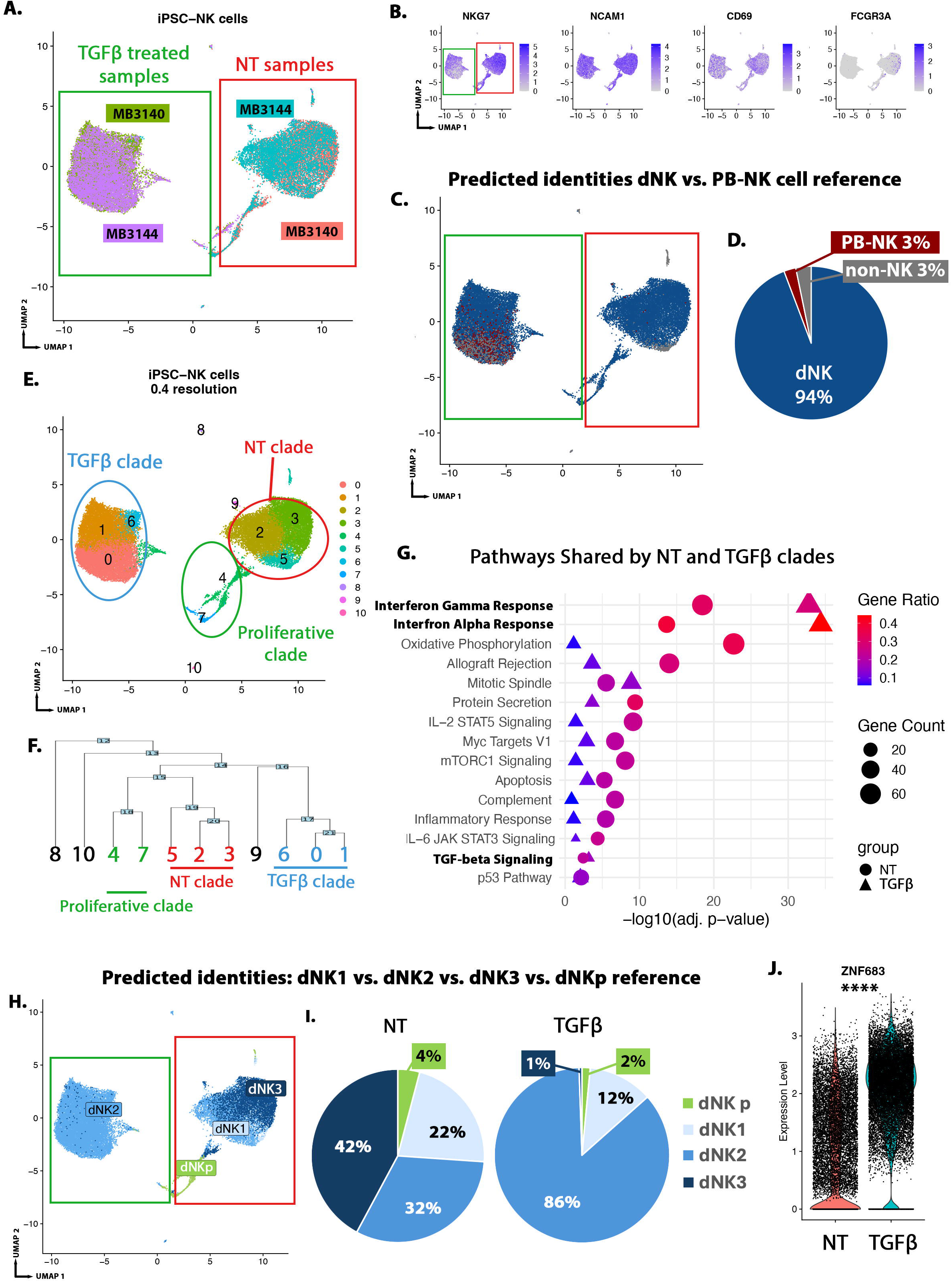
iPSC-NK cells transcriptionally resemble dNK cells, with TGFβ promoting a dNK2 phenotype. 2 iPSC lines (MB3140 and MB3144 lines) differentiated with and without TGFβ were subjected to scRNA-seq, resulting in 4 samples with a total of 33,830 cells after filtering for analysis (MB3140_NT: 9147 cells; MB3144_NT: 6628 cells; MB3144_TGFβ: 9152 cells; MB3140_TGFβ: 10,031 cells). **A)** UMAP showing no treatment (NT, no TGFβ) and TGFβ iPSC-NK cells derived from MB3140 and MB3144 iPSC lines. Unsupervised clustering resulted in separation by treatment group with TGFβ-treated cells on the left and NT on the right. **B)** FeaturePlots showing expression of dNK marker genes (*NKG7*, *NCAM1*, *CD69*) and absence of PB-NK marker gene *FCGR3A*, which encodes CD16. **C)** UMAP showing predicted identities for iPSC-NK, using all cells from Vento-Tormo et al., 2018^22^ with dNK (comprised of dNK1, dNK2, dNK3, and dNKp) and PB-NK (comprised of NK CD16+ and NK CD16-) as reference. **D)** Pie chart quantification of data in **(C)**. **E)** UMAP showing clustering of iPSC-dNK at 0.4 resolution. **F)** Dendrogram showing phylogenetic analysis of aggregate gene expression of iPSC-dNK clusters; TGFβ clade in blue, NT clade in red, Proliferative clade in green. **G)** Quantification of pathways significantly enriched in both NT and TGFβ clades using MSigDB Hallmark 2020 pathway analysis. There are two values per pathway, one each from NT clade-enriched DEGs (circles) and TGFβ clade-enriched DEGs (triangles). **H)** UMAP showing all predicted identities for iPSC-dNK cells, using the dNK cell subtypes (dNK1, dNK2, dNK3, dNKp) from Vento-Tormo, 2018^22^ as reference. **I)** Pie chart quantification of data in **(H)**. **J)** *ZNF683*, dNK2 marker gene, has significantly higher expression in TGFβ iPSC-dNK in comparison to NT iPSC-dNK. Statistical analysis was performed by Wilcox Rank Sum test using R. ****p < 0.0001 Alt text: UMAPs showing NK cell marker expression by iPSC-NK. UMAPs and Pie chart showing dNK cell identify of 94% of iPSC-NK. UMAP and dendrogram showing clustering of iPSC-dNK by treatment group. Graph showing gene enrichment in dNK-associated pathways by iPSC-dNK, with further enrichment upon TGFbeta treatment. UMAP and pie chart showing iPSC-dNK resemble all dNK subtypes with increased proportion of dNK2 upon TGFbeta treatment. Violin Plot showing higher expression of dNK2-marker gene ZNF683 in TGFbeta iPSC-dNK

We next investigated iPSC-dNK heterogeneity using unsupervised and supervised analyses. Unsupervised clustering of iPSC-dNK resulted in 11 clusters, with clusters 0-7 representing the majority (99%) of cells (**Figure 3E**; **Supplementary Figure 4B**). Hierarchical clustering grouped clusters 0, 1, and 6 into one clade, comprised of TGFβ iPSC-dNK (**Figure 3F**), expressing TGFβ-inducible genes, *JUN* and *SMAD7* (**Supplementary Figure 4C**). Clusters 2, 3, and 5 were grouped into another clade, comprised of no treatment (NT) iPSC-dNK (**Figure 3F**). Clusters 4 and 7 were grouped into a third clade, comprised of G2M and S phase cells expressing proliferation genes (*MKI67*, *PCNA*). (**Supplementary Figure 4D,E**). Finally, clusters 8, 9, and 10, comprising a small minority (0.5%) of cells, mapped primarily to non-NK populations (**Figure 3C**).

We identified marker genes for each cluster and continued with gene enrichment analysis of each of the major clades (**Supplementary File 2**). Both the NT and TGFβ clades were enriched for many pathways involved in NK cell function such as Protein Secretion, IL6 JAK/STAT3 signaling, and IL-2/STAT5 signaling (**Supplementary File 3**, **Figure 3G**).^51–53^ We also found an enrichment for dNK-specific signaling pathways, Interferon Alpha and Gamma Response, and TGF-beta signaling (**Figure 3G**, **Supplementary Figure 3B**). Interestingly, all three of these dNK-specific pathways were more significantly enriched within the TGFβ clade (**Figure 3G**, triangle). Additionally, we found that there was an enrichment of genes involved in both “TGF-beta signaling” and “PI3K/AKT pathway” in the TGFβ clade, suggesting that our TGFβ treatment was acting via both canonical and non-canonical pathways (**Supplementary File 3**).^54^ As expected, DEGs in the proliferating clade were enriched for cell cycle control (**Supplementary File 3**). Notably, there was enrichment for “Lymphoid cells of the Placenta” in both NT and TGFβ clade enriched DEGs (**Supplementary Tables 1-3**), consistent with the dNK identity predicted by reference mapping. However, pathway analysis identified differential activation of immune signaling pathways between NT and TGFβ clades suggestive of more nuanced differences between the iPSC-dNK. iPSC-dNK showed dNK-associated gene expression that was further enriched by TGFβ.

To compare the heterogeneity of the iPSC-dNK to that of dNK *in vivo,* we extracted the dNK subclusters (dNK1, dNK2, dNK3, dNKp) from Vento-Tormo et al.,^22^ to use as a reference for iPSC-dNK (**Supplementary Figure 4F**). We found the majority of cells (42%) in the NT clade mapped to dNK3, with the next largest population of cells (32%) mapping to dNK2, followed by dNK1 (22%), while the TGFβ clade found 86% of cells mapping to dNK2, 12% to dNK1, and only 1% to dNK3 (**Figure 3H,I, Supplementary Figure 4G**). Further, the iPSC-dNK expressed dNK, but not PB-NK, marker genes, and showed an enrichment of dNK2 marker gene *ZNF683* within the TGFβ clade (**Figure 3J, Supplementary Figure 4A**).

In summary, scRNA-seq analysis confirmed the dNK identity of iPSC-dNK. We identified heterogeneity within the iPSC-dNK, representing relevant dNK subtypes, finding that TGFβ further enriches for dNK signaling pathways and alters the proportion of dNK subtypes in favor of dNK2.

### iPSC-dNK functionally resemble dNK

With the dNK identity of the iPSC-dNK established, we set out to analyze their function. dNK from early and late gestation express cytolytic proteins.^13^ We found that NT and TGFβ iPSC-dNK expressed *GNLY*, *PRF1*, and *GZMB* (**Supplementary Figure 5A**) by scRNA-seq. We also found protein expression, albeit at low levels, of PFN (encoded by *PRF1*) and the first trimester dNK-specific 9kD GNLY isoform in both NT and TGFβ iPSC-dNK (**Supplementary Figure 5B-C**).^13^

First trimester dNK, not PB-NK, secrete IL-6, VEGF, and PLGF in the absence of stimulation.^15,16^ However, it is not known if term or iPSC-dNK share this function. We applied aptamer-based proteomic profiling, using the SomaScan Assay 7K panel (See **Methods**) to conditioned media from purified NK cells from peripheral blood, first trimester decidua, and term BP and CAM, and compared them to those of NT iPSC-dNK and TGFβ iPSC-dNK (**Figure 4A**). We found 4,938 proteins secreted by primary cells at levels above media-only control, of which 1,456 were found in first trimester, BP, or CAM dNK, but not PB-NK (**Figure 4B**); of these 1,456 proteins, 570 were shared by all 3 primary dNK (**Supplementary Figure 6**). We found iPSC-dNK secreted proteins had substantial overlap with those from primary dNK, and not PB-NK (**Figure 4C**). TGFβ induced secretion of a markedly higher number of dNK-specific proteins, compared to NT iPSC-dNK (217 in TGFβ iPSC-dNK, vs. only 28 in NT iPSC-dNK) (**Figure 4C**).

**Figure 4.**
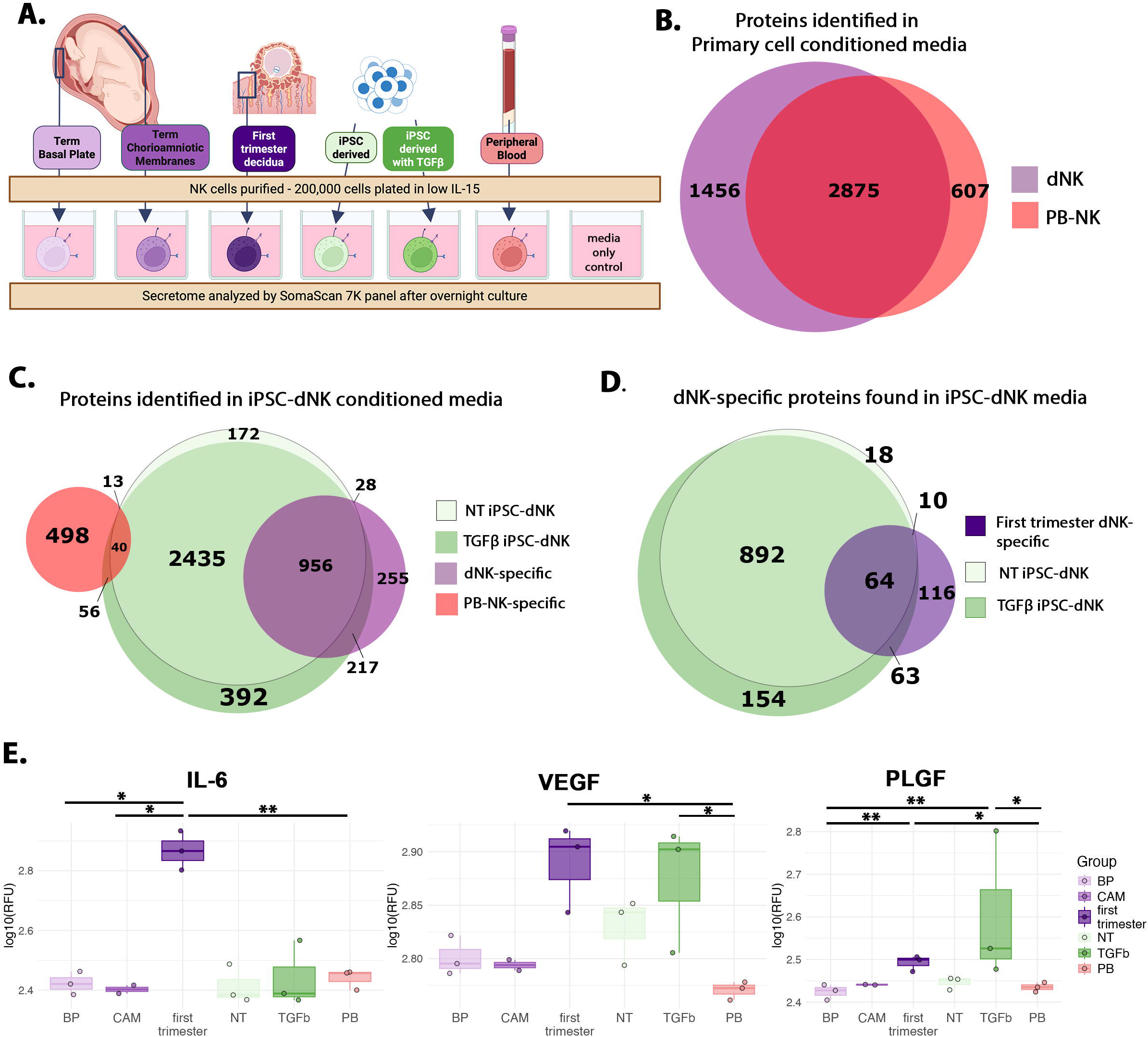
iPSC-dNK secrete proteins similar to primary dNK. **A)** Schematic of experimental design for aptamer-based proteomic profiling. **B)** Venn Diagram showing 2,875 proteins identified in conditioned media of first trimester, term BP-, and term CAM-dNK, as well as PB-NK; 1,456 proteins found only in primary dNK media (dNK-specific); and 607 proteins found only in PB-NK media (PB-NK-specific). **C)** Venn Diagram showing overlap of proteins identified in secretome of NT and TGFβ iPSC-dNK with dNK-specific (1,456) and PB-NK-specific (607) proteins; significantly higher overlap was noted with the dNK-specific proteins. **D)** Venn Diagram showing overlap of proteins identified in secretome of NT vs. TGFβ iPSC-dNK with first trimester dNK-specific (253) proteins; significantly more overlap was noted with TGFβ iPSC-dNK proteins. **E)** Box plots showing median expression as well as individual log10RFU values for each sample. Proteins known to be enriched in dNK (over PB-NK) are shown: IL-6 (seq.2573.20), VEGF (seq.2597.8), and PLGF (seq.2330.2). Term BP n= 3, Term CAM n=2, First Trimester decidua n =3, NT iPSC-dNK n = 3, TGFβ iPSC-dNK n=3, PB-NK n=3. Dunn’s test for multiple comparisons, using R, is shown; *p<0.05, **p<0.01. Alt text: Schematic illustrating all NK cells analyzed by aptamer-based proteomic profiling. Venn diagrams showing primary dNK secrete more proteins than PB-NK. Venn diagram showing iPSC-dNK secreted more dNK-specific proteins than PB-NK-specific proteins. Venn diagram showing TGFbeta iPSC-dNK express more first trimester dNK specific proteins than NT iPSC-dNK. Box and whisker plots showing higher levels of IL-6, VEGF, and PLGF proteins in first trimester dNK and TGFbeta iPSC-dNK.

Transcriptionally, TGFβ iPSC-dNK were enriched for dNK2, the most represented subtype in first trimester dNK (**Figure 1B**). We therefore asked if the secretory function of TGFβ iPSC-dNK reflected that of first trimester dNK. Of the 253 first trimester dNK-specific proteins (**Supplementary Figure 6A**), 127 were identified in TGFβ-treated iPSC-dNK secretome while only 74 were identified in NT iPSC-dNK (**Figure 4D**). We saw the same enrichment when we considered all 3,683 proteins secreted by first trimester dNK, with 3,231 also secreted by TGFβ iPSC-dNK (**Supplementary Figure 6B**). In agreement with prior literature, we found levels of IL-6, VEGF, and PLGF were higher by 2.5-fold, 2.7-fold, or 2.1-fold, respectively, in first trimester dNK, compared to PB-NK (**Figure 4E**). We also report, for the first time, that high IL-6 and PLGF are in fact specific to first trimester dNK, as their levels are lower in BP- and CAM-dNK secretomes, which are more similar to that of PB-NK (**Figure 4E**). Interestingly, we found TGFβ iPSC-dNK secrete higher levels of VEGF and PLGF, comparable to first trimester dNK (**Figure 4E**).

In comparison to first trimester dNK, primary term dNK and PB-NK have similarly higher degranulation (CD107a) in response to PMA/I stimulation. However, compared to PB-NK, stimulated first trimester and term dNK have lower production of cytokines (IFNγ, TNFα).^13^ GM-CSF, on the other hand, has a reciprocal response, with higher induction in first trimester dNK than PB-NK. ^23^ Within first trimester dNK subtypes, in comparison to dNK2, dNK3 responded with higher levels of CD107a, GM-CSF, and IFNγ.^23,24^ We found iPSC-dNK to be responsive to PMA/I stimulation, inducing CD107a, IFNγ, TNFα, and GM-CSF, with decreased production of TNFα and GM-CSF in TGFβ-treated cells (**Figure 5A-D**).

**Figure 5.**
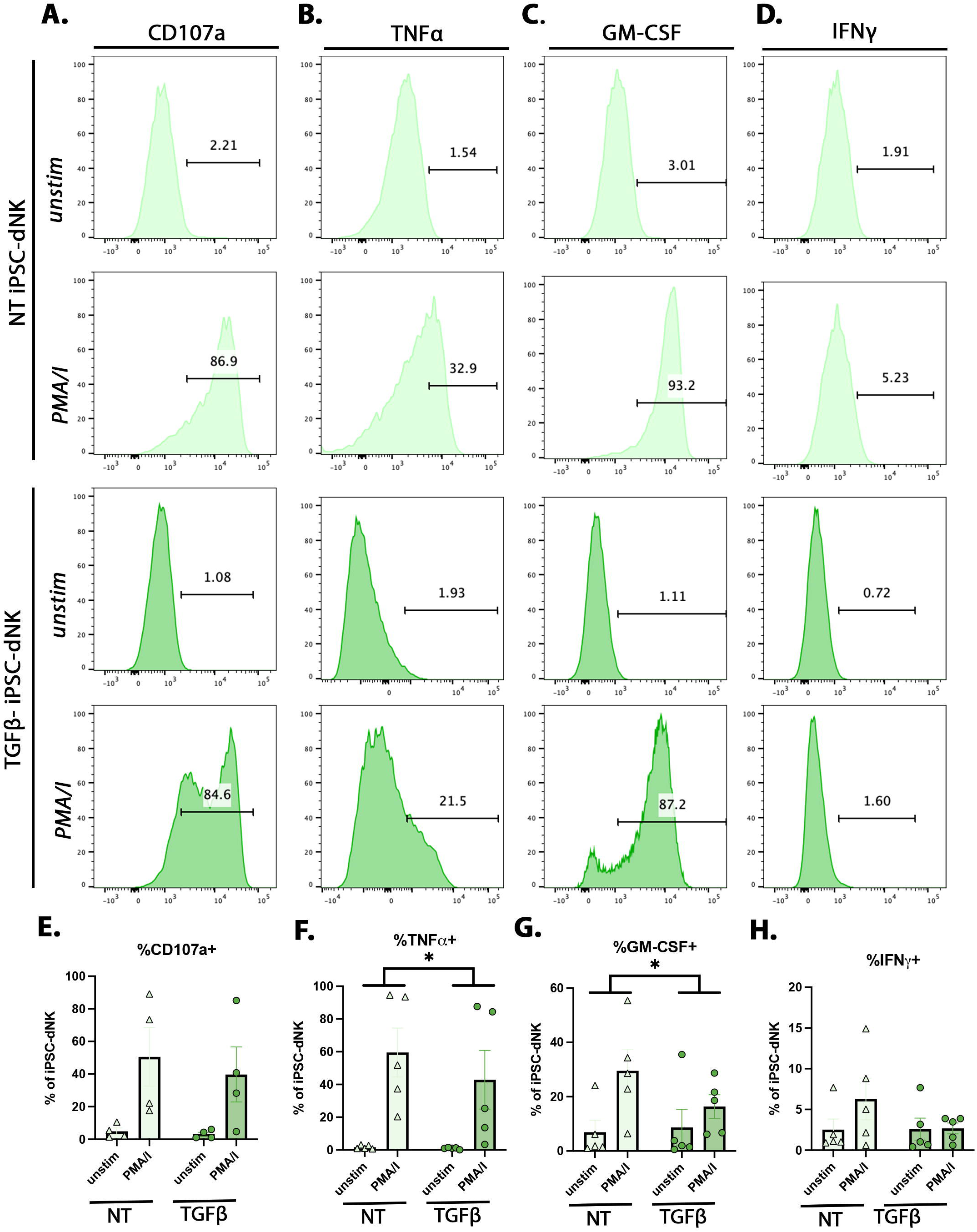
iPSC-dNK response to stimulation is modulated by TGFβ. Representative FACS plots for **A)** CD107a **B)** TNFα **C)** GM-CSF, and **D)** IFNγ of NT and TGFβ iPSC-dNK in the presence or absence (unstim) of PMA/I treatment. Quantification of stimulation markers **E)** CD107a **F)** TNFα **G)** GM-CSF, and **H)** IFNγ, as a percent of expression of dNK, as measured by flow cytometry. Statistical testing was performed on PMA/I – unstim of n=4 independent experiments are iPSC-dNK derived from MB3140 and MB3144 iPSC lines. Data in **E** through **H** are represented as mean +/- standard error, by paired t-test using GraphPad Prism; *p<0.05. Alt text: Histograms and bar graph quantification showing iPSC-dNK increase CD107a, TNFalpha, GM-CSF, and IFNgamma in response to PMA/I stimulation. Statistical analysis shown on bar graphs show decreased TNFalpha and GM-CSF by TGFbeta iPSC-dNK

Finally, we evaluated the cytotoxicity of NT and TGFβ iPSC-dNK relative to term CAM-and BP-dNK, first trimester dNK, PB-NK, and NK-92 cells, against JEG3 choriocarcinoma target cells, at 1:1, 2:1, and 5:1 effector-target (E:T) ratios (**Figure 6A**). In alignment with the current literature, we found that first trimester dNK trend towards inducing apoptosis in target cells at reduced levels relative to NK-92 and PB-NK (**Figure 6B-C**). Interestingly we found that increasing numbers of BP- or CAM-dNK do not induce increased apoptosis in target cells (**Figure 6B-C**). Finally, we found that NT and TGFβ iPSC-dNK were able to kill target cells (**Figure 6B-C**), demonstrating functional maturity, similar to primary first trimester dNK and PB-NK.

**Figure 6.**
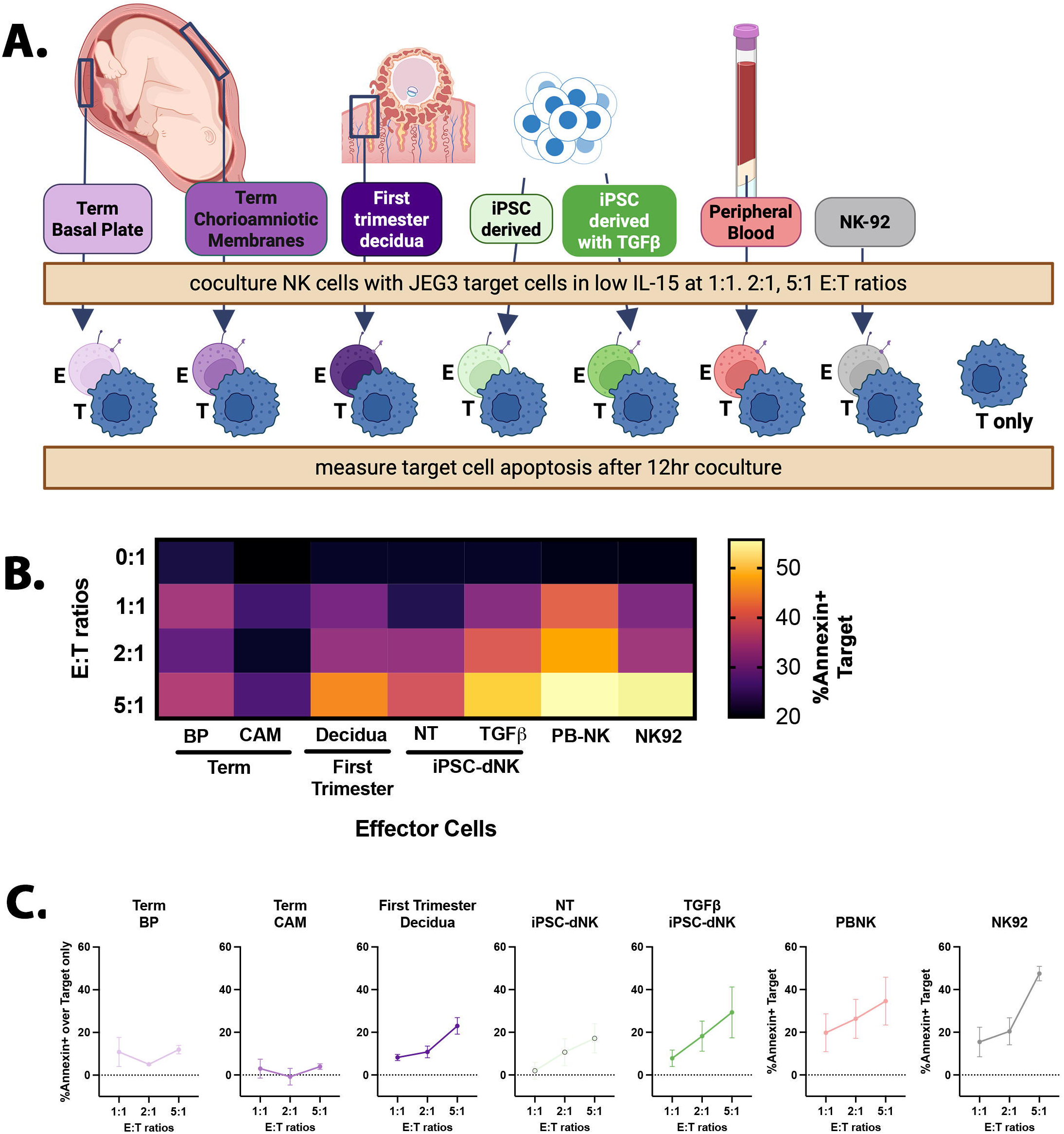
iPSC-dNK have cytotoxic activity. **A)** Schematic of all NK effector cells (E) tested for cytotoxicity against JEG3 choriocarcinoma target cells (T). **B)** Heatmap representation of mean expression of Annexin+ target cells at the indicated effector-to-target (E:T) ratios. **C)** Percent Annexin+ (apoptotic) EGFR^+^CD45^-^ target cells subtracted from baseline (JEG3-only) apoptosis, after coculture with different effector cells at the indicated E:T ratios. Term BP, n=2 biological replicates; Term CAM, n=3 biological replicates; First trimester decidua, n=3 biological replicates, n = 4 technical replicates; NT iPSC-dNK n = 3 biological replicates, n=4 technical replicates; TGFβ iPSC-dNK, n=3 biological replicates, n=4 technical replicates; PB-NK, n=3 biological replicates, n=4 technical replicates; NK92, n=2. Three independent experiments were performed. JEG3 (Target cell)-only control Annexin expression included in each experiment and subtracted from all samples for analysis. Statistical analysis was performed using Two-way ANOVA with Sidak’s test for multiple comparisons using GraphPad Prism; Data is shown as mean ± standard error. Effector cell is found to be a statically significant source of variation with a ***p-value=0.0007. E:T ratio is found to be a statically significant source of variation with a **p-value=0.0022. Sidak’s test for multiple comparisons finds Term CAM dNK vs. PBNK ** adj.p-value=0.0015; Term CAM dNK vs. NK92 ** adj.p-value=0.0095; NT iPSC-dNK vs. PBNK *adj.p-value=0.05. All other pairwise comparisons have p-values >0.05. Adj.p-values for all pairwise comparisons can be found in **Supplementary Table 5**. Alt text: Schematic illustrating all NK cell function analyzed by Killing Assay. Heat map showing first trimester dNK, NT iPSC-dNK, TGFbeta iPSC-dNK, PB-NK, and NK-92 effector cells kill JEG3 target cells, and this increases with increasing E:T ratios. Line graphs showing the average level of apoptosis above baseline cell death in target cells.

In summary, we found that, overall iPSC-dNK are functionally most similar to primary dNK, and that TGFβ iPSC-dNK have increased secretion of first trimester dNK-specific proteins and reduced stimulation response, supportive of a first trimester dNK (dNK2-dominant) phenotype.

## Discussion

dNK are a key cell type during pregnancy, playing vital roles from implantation in early gestation, to continued regulation of EVT function at the basal plate (BP), and regulation of membrane rupture in chorioamniotic membranes (CAM) during labor. These various functions may be mediated by different dNK subtypes.^22–24^ We found dNK subtype compositions between first trimester decidua and term BP and CAM reflected this change in function. We also present novel analysis of NK cell function in our study, and report the secretome of first trimester dNK, term BP-and CAM-dNK, and PB-NK, as well as the first characterization of term BP- and CAM-dNK cytotoxicity. We used these primary cell data, in combination with published literature to develop a rigorous dNK scorecard, against which to benchmark newly developed models (**Figure 7**). To this end, we report a robust, reproducible, and tunable protocol for induction of functional dNK from iPSC, as measured by surface markers, transcriptome, secretome, and functional activation.

In agreement with previous literature, we found term dNK show greater expression of activating receptors, compared to first trimester dNK. We also identified a high proportion of high-effector function dNK3 at term, increased from first trimester, suggesting this increase in dNK3 is mediating the increased activating receptors, decreased inhibitory receptors, and increased PMA/I response observed in term dNK. However, while we found BP-dNK to have the greatest proportion of dNK3 based on scRNA-seq analysis, percent positivity for CD103 (a dNK3 marker) was not significantly increased, relative to first trimester or term-CAM dNK, suggesting discordance between transcriptomic and proteomic subtype definitions. Previous studies have found that term dNK show decreased expression of some cytolytic molecules (GNLY, 9kD GNLY), but not others (PFN, GZMB), relative to first trimester dNK. In comparison with first trimester dNK, term dNK degranulation is higher in response to PMA/I and K562 target cells, but lower in response to HCMV-infected decidual stromal cells.^13^ As such, it is unclear whether term dNK possess the capacity to kill a target cell and how this capacity compares to first trimester dNK and PB-NK. We found that term BP- and CAM-dNK did not induce apoptosis of JEG3 cells with increasing E:T ratios, while first trimester dNK did, albeit at levels below PB-NK and NK-92. Our study provides new information on how dNK are functionally changed throughout gestation and decidual locations reflecting the dynamic nature of the maternal-placental interface while highlighting the need for more comprehensive definitions for dNK subtypes.

**Figure 7.**
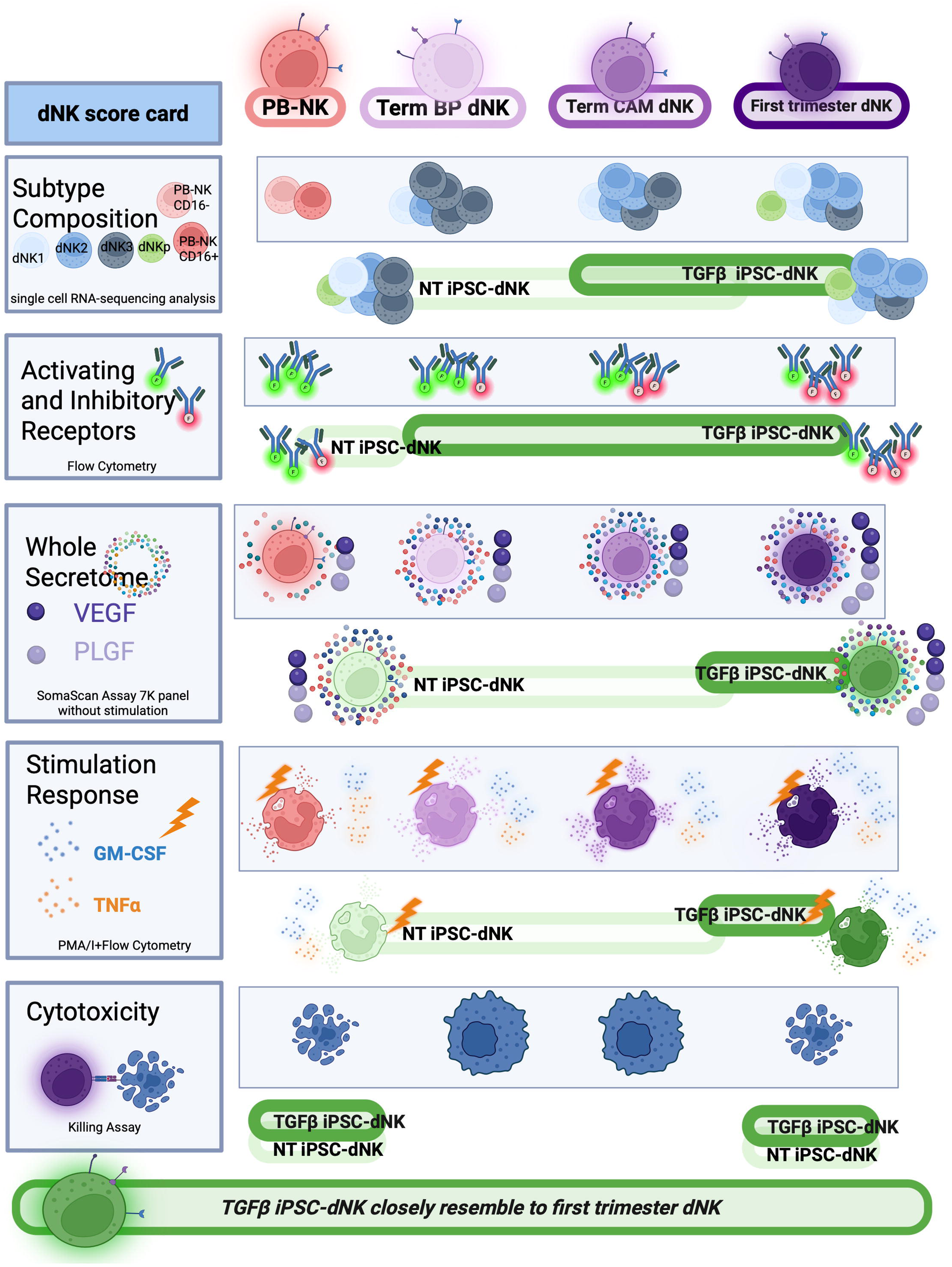
TGFβ iPSC-dNK most closely resemble first trimester dNK. Summary schematic of key features of NK cells as measured in PB-NK and dNK across gestation and different placental locations as well as the results of benchmarking NT and TGFβ iPSC-dNK against these definitions. Alt text: Schematic illustrating all assays used to define dNK identity and the results of assessing NT and TGFbeta iPSC-dNK against these criteria.

dNK heterogeneity is an emerging area of study. While variegated expression of cell surface markers by dNK have long been observed, the field has only recently begun defining dNK subtypes as scRNA-seq technologies and analysis tools have improved. As such, transcriptional definitions are most universally accepted, while cell surface markers and functional outputs associated with dNK1, dNK2, and dNK3 are still being defined. While this limits the conclusions we can draw from our own data, our study begins to bring the transcriptomic, flow cytometric, and functional definitions of dNK together, using primary and iPSC-derived dNK. Importantly, we expect our iPSC-derived model to accelerate research in this space.

Studies of term dNK have shown no differences in protein expression or function based on localization in BP vs. CAM. However, term dNK from the two compartments do show gene expression differences.^13^ Single-cell analyses have pointed to a relative increase in dNK in CAM, but not BP, in placentas of patients with preterm labor.^49,55^ Further, labor-associated differential interactions have been identified in full term BP-dNK and CAM-dNK.^56^ Our reanalysis of these data identified similar differences between BP- and CAM-dNK.^49^ Notably, we found that ∼50% of the dNK in CAM are dNK2, similar to first trimester, while the dNK2 is reduced (to 23%) in BP. dNK3 dominance over dNK2 in BP and dNK2 over dNK3 in CAM was consistent between term with and without labor (see **Supplementary Figure 1J**). Interestingly, we observed increased dNK3 in CAM in placentas with labor (term or preterm) but decreased dNK3 in BP of placentas from patients with preterm labor. Due to the small sample size (n=3 per labor group), we conclude that these observations warrant follow up studies on dNK subtypes in both BP and CAM in the context of labor, particularly preterm labor.

We found the proportion of dNK2 (scRNA-seq) and CD9^+^CD56^bright^CD16^-^ dNK (flow cytometry) were similar between first trimester decidua and term CAM; meanwhile, term BP showed decreased dNK2 based on scRNA-seq and decreased CD9^+^CD56^bright^CD16^-^ by flow cytometry. While the frequencies of CD9^+^ dNK (∼90%) and dNK2 (∼50%) are not the same within a single tissue, these data suggest CD9 could be used, in combination with other markers, to define dNK2. We add further support for this relationship using our newly-developed differentiation protocol for iPSC-dNK. TGFβ treatment resulted in selection against dNK3, in favor of dNK2, based on transcriptome, increased CD9 by flow cytometry, as well as reduced GM-CSF secretion by PMA/I stimluation. These data from both primary and iPSC-dNK suggest that CD9, in combination with other markers, may be useful to identify a dNK2 transcriptional phenotype; however, additional experiments are needed to validate this concept.

Our study is the first application of untargeted aptamer-based proteomic profiling of secretome from primary NK cells, including first trimester and term dNK and PB-NK. We applied this method to iPSC-dNK and functionally compared them to primary dNK. Of the more than 4,000 proteins identified in the secretomes of primary NK cells, we found approximately 1/3 to be dNK-specific. We also found similarities in dNK across gestation – with significant overlap between first trimester and term BP dNK - as well gestational age-related changes, such as reduced secretion of proangiogenic proteins at term (**Supplementary Figure 6A**). TGFβ treatment is typically shown to suppress NK cell activation and function.^34,35,57^ Notably, however, TGFβ treatment of PB-NK alone does not induce VEGF secretion.^58^ We found that TGFβ iPSC-dNK secrete VEGF at levels significantly higher than PB-NK (**Figure 4E**), supportive of TGFβ promoting differentiation to dNK. Interestingly, we observed no enrichment of *VEGF* transcript by any dNK subtype in first trimester decidua (data not shown); however, VEGF levels were similar in TGFβ iPSC-dNK and first trimester dNK secretomes, suggesting this could be a functional signature of dNK2. However, follow up studies are required to validate this interpretation.

Historically, iPSC-derived cells have been found to be immature and not fully functional.^36^ However, this is not the case for iPSC-derived NK cells, as they have been used *in vivo* and in the clinic.^43,44,59^ While short term exposure to TGFβ is associated NK cell suppression,^43^ long-term exposure to TGFβ during NK activation, known as “TGFβ imprinting,” is associated with heightened cytokine secretion, degranulation, and cytotoxicity.^60^ We find that NT and TGFβ iPSC-dNK are functionally mature, as shown by their secretion of cytokines and angiogenic proteins, as well as their cytotoxicity against target cells. Further, TGFβ iPSC-dNK had increased secretion of angiogenic proteins and a trend towards higher induction of apoptosis in target cells, perhaps consistent with the dNK2 phenotype, which reportedly show moderate cytokine induction and degranulation in response to PMA/I stimulation;^23,24^ however cytolytic protein expression and cytotoxicity of dNK subtypes have not been evaluated. Overall, our findings emphasize the need for integrated transcriptomic and functional definitions of dNK subsets.

A major limitation of our model system is that our iPSC-dNK are derived and cultured in the absence of *in vivo* spatial, endocrine, and cellular cues. Previous scRNA-seq analyses have identified potential interaction partners between dNK and other decidual cell type, such as EVT and dESF.^22^ However, due to limitations in access and cell yield, it is difficult to use primary cells to develop more complex multi-cell type 3-dimensional models to study such *in vivo* cues and validate these hypothesized interactions. We have previously generated maternal dESF and placental EVT from iPSC and here established a protocol for differentiation of dNK from iPSC.^38,61^ Our iPSC-based models can overcome limitations, such as cell yield which introduces allogenic incompatibilities, in particular between maternal dESF and dNK, in cocultures. Further, these models offer technical advantages for mechanistic studies using gene editing.^43,44^ In our NT and TGFβ iPSC-dNK, we observed expression of *CCL5* and *XCL1*, known partners of *CCR1* and *XCR1*, respectively, expressed by EVT, macrophages, and dendritic cells.^22^ Furthermore, we can induce specific receptors and ligands by modulating expansion phase media composition. Specifically, NT iPSC-dNK showed high *XCL2* and *KLRB1*, partners for *XCR1* and *CLEC2D*, respectively, expressed by EVT, dESF, and dendritic cells; conversely, TGFβ iPSC-dNK showed high *CXCR4* and *CSF1*, partners for *CXCL12* and *CSFR1*, respectively, expressed by dESF and macrophages.^22^ The abundant cell yield and the ability to control receptor expression without antibody purification - and thus interference - will enable studies of specific interactions affecting placentation and pregnancy outcomes between dNK and other cell types.

A second limitation of this study is that we cannot rule out transcriptional similarity of our iPSC-dNK to tissue-resident (tr)-NK of other tissues. Indeed, this would be an interesting line of investigation, as dNK share expression of CD69 and CD103 with trNK cells, found in the lungs.^28,63,64^ scRNA-seq of human blood, tonsils, lung, and intestinal mucosa clustered NK cells into 4 broad categories: CD56^Bright^ and CD56^Dim^, both which can be found in tissues and blood, and PRDM1^+^ ILC1 and ZNF683^+^ NK, found only in tissue. ^63,64^ The authors compared these NK subtypes to dNK and found dNK1 was most similar to intestinal intraepithelial PRDM1^+^ ILC1, while dNK2 and dNK3 most closely resembled ZNF683^+^ NK found in lung and tonsil. We identified *ZNF683* as a dNK2 marker which can be enriched for in iPSC-dNK by TGFβ. These data suggest that follow up analyses of TGFβ iPSC-dNK may find resemblance to lung and tonsil ZNF683^+^ NK, broadening the applications of our protocol.

Just as iPSC-NK are being developed as cellular therapies for cancer treatment, it is paramount that we also develop these technologies into therapeutics for reproductive conditions. Our protocol offers a first step toward development of such therapeutics, and hopefully, the prevention of adverse pregnancy outcomes.

## Resource Availability

### Materials availability

This study did not generate new unique reagents. Requests for further information, reagents, and human pluripotent stem cell lines used in this study should be directed and will be fulfilled by the Lead contact, Mana Parast.

### Data and code availability

- Single-cell RNA-seq data are deposited at GEO and publicly available as of the date of publication. Accession numbers will be listed in the key resources table (GSE302146).
- This paper does not report original code. The final code used for data processing and analysis will be deposited at Zenodo and made publicly available as of the date of publication. DOIs are listed in the key resources table.
- Any additional information required to reanalyze the data reported in this paper is available from the corresponding author upon request.

## Acknowledgements and Funding

The authors are grateful to all patients who donated tissues for this research. This work was funded by the National Institutes of Health (NIH) (R01-HD102639 to M.M.P. and J.D.B.; R00-HD091452 to M.H.), NIH/NCATS 2UL1TR001442-08 (CTSA), as well as faculty-generated discretionary funds. V.C.C. was supported by F32-HD108944 and T32-HD007203 to the University of California, San Diego. M.H. was also supported by the UCSD Academic Senate grant RG106839. J.D.B. was also supported by The Hartwell Foundation. C.D. was supported by the California Institute for Regenerative Medicine Bridges Grant EDUC2-08376 awarded to San Diego State University. J.J. supported by the California Institute for Regenerative Medicine Bridges Grant EDU-1261 awarded to California State University San Marcos. This publication includes data generated at the UC San Diego IGM Genomics Center utilizing an Illumina NovaSeq 6000 that was purchased with funding from a National Institutes of Health SIG grant (#S10 OD026929). Computational analysis was performed on the Extreme Science and Engineering Discovery Environment (XSEDE) Expanse at SDSC, which is supported by National Science Foundation grant number ACI-1548562 (allocation ID: BIO220095). Graphical abstract and schematics in Figure 1, Figure 2, Figure 4, Figure 6, Figure 7, and Supplementary Figure 2 were created in BioRender. Cheung, V. (2026) https://BioRender.com/pxhkijv

## Disclaimers

V.C.C., J.J., C.D., H.A., C.C., J.S., M.F., M.M., K.F., R.E.M., L.S.C., D.P., M.H., J.D.B, and M.M.P declare no competing interests. D.S.K. is a co-founder and advisor to Shoreline Biosciences and has an equity interest in the company. D.S.K. also consults Therabest and RedC Bio for which he receives income and/or equity. Studies in this work are not related to the work of those companies. The terms of these arrangements have been reviewed and approved by the University of California, San Diego, in accordance with its conflict-of-interest policies.

---

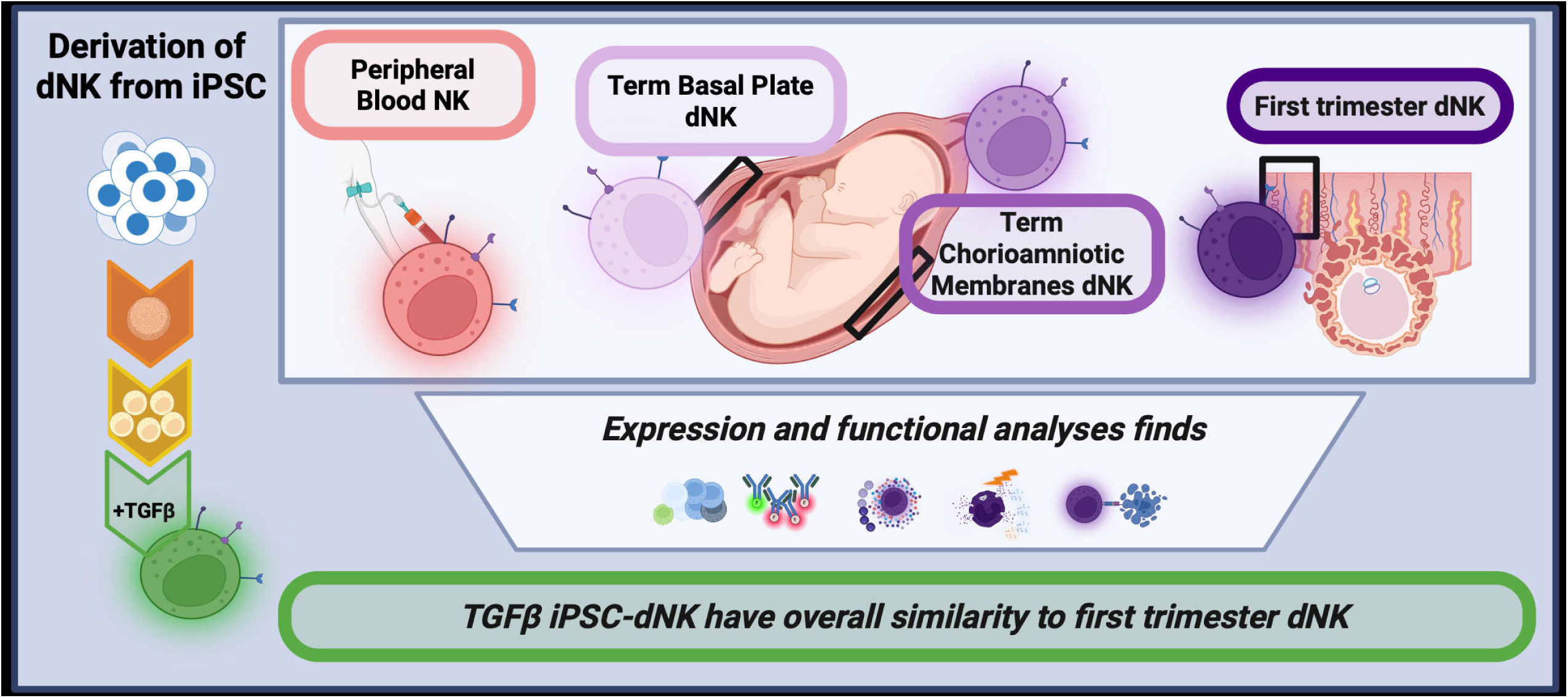

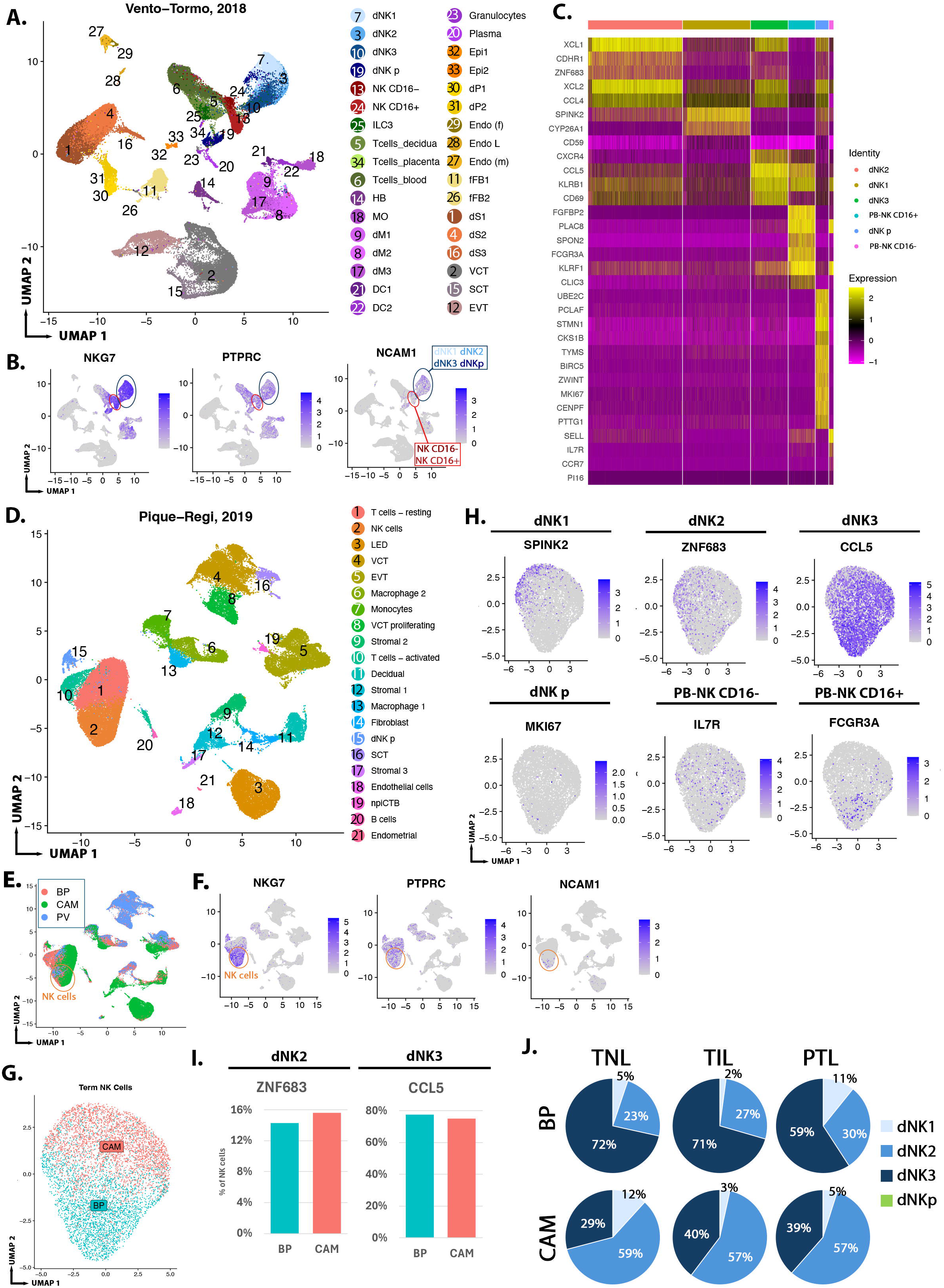

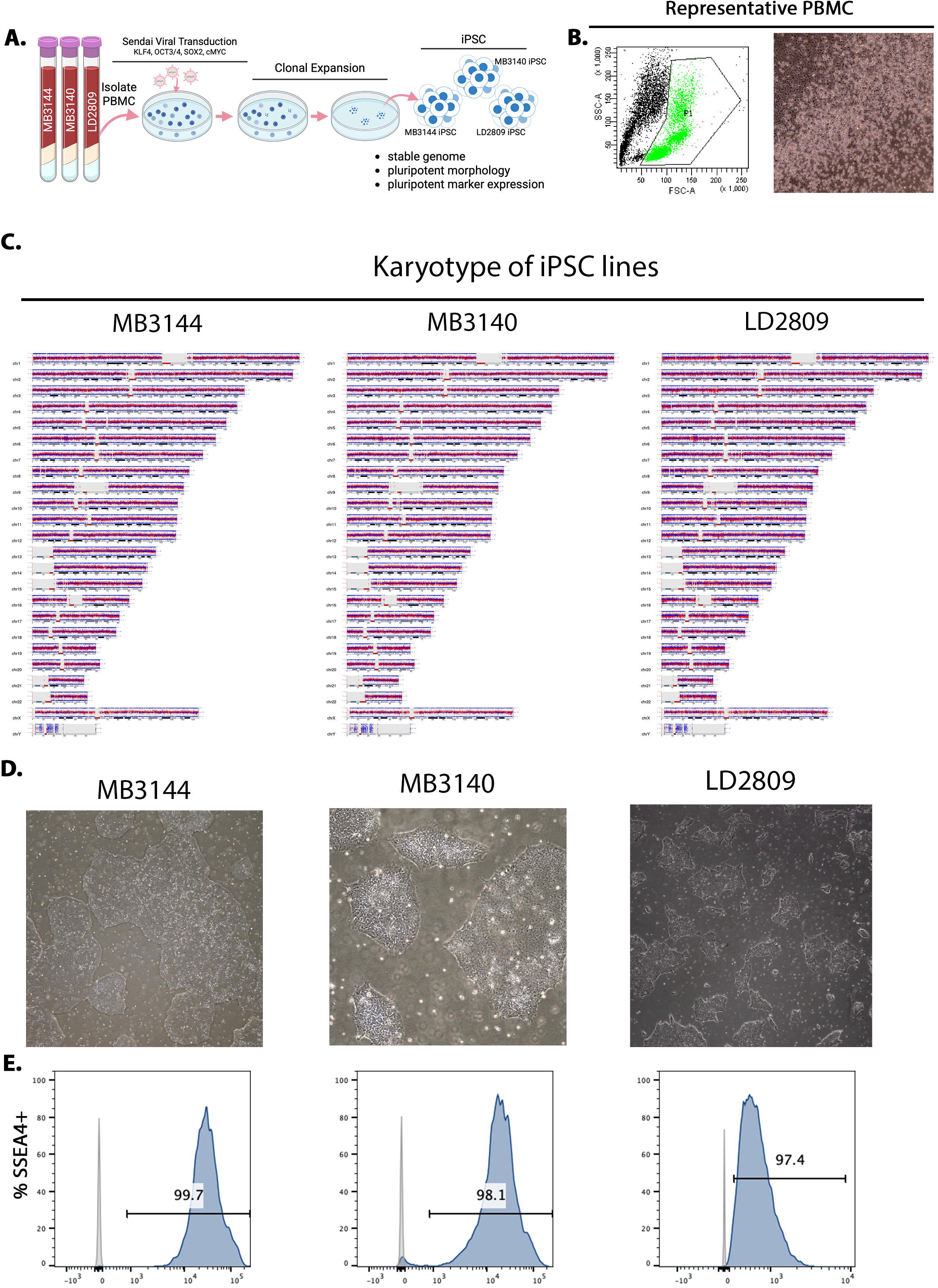

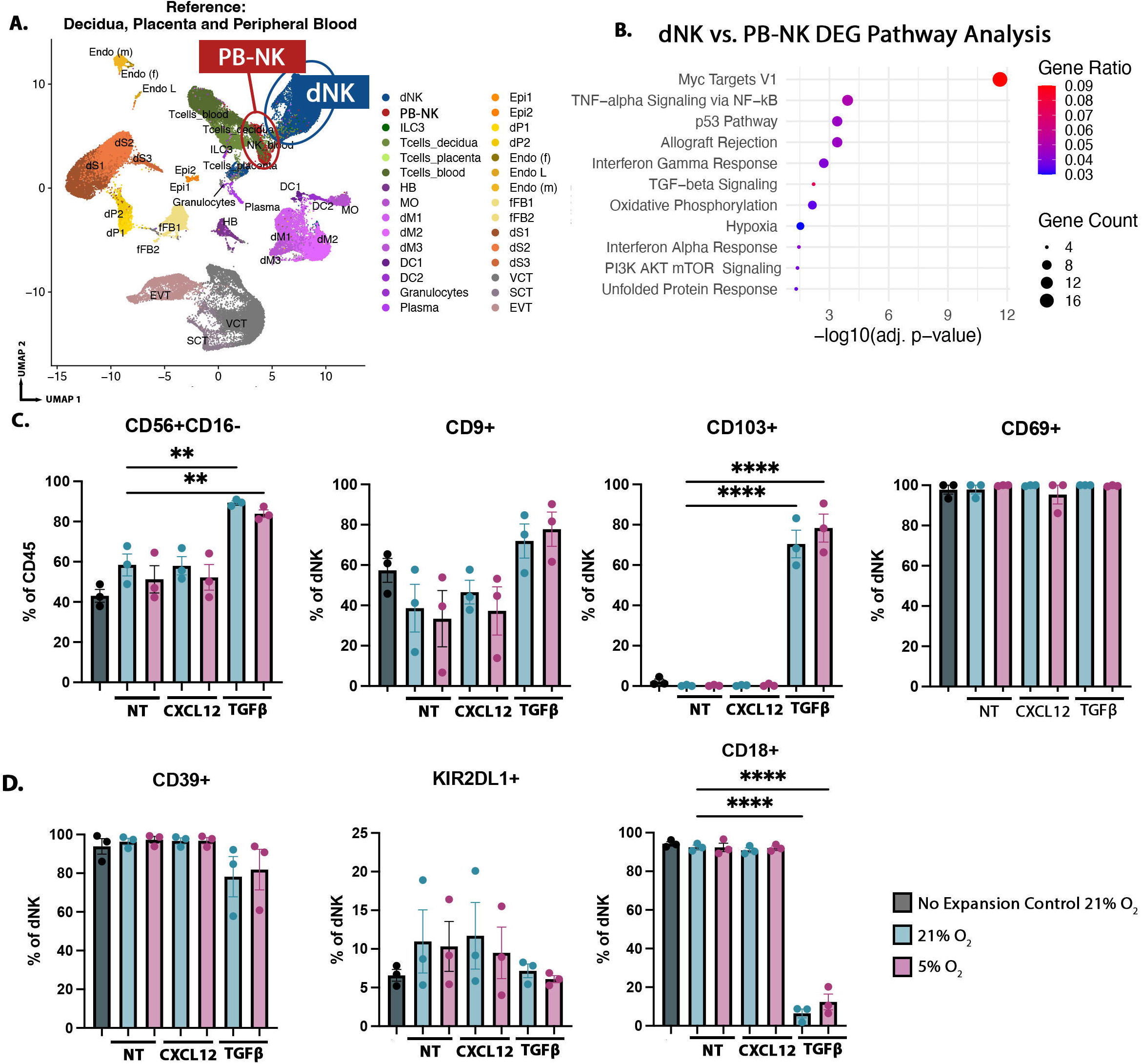

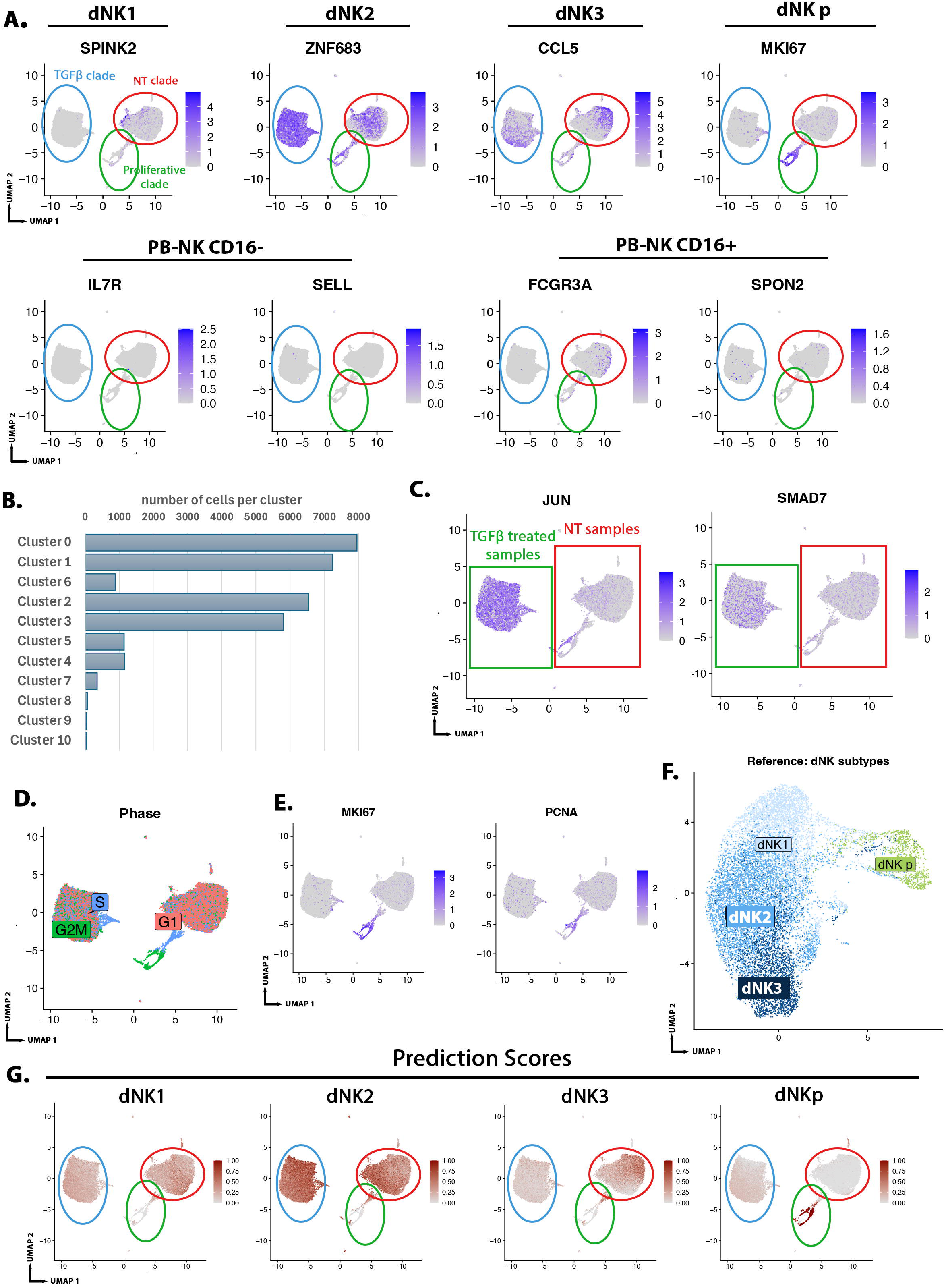

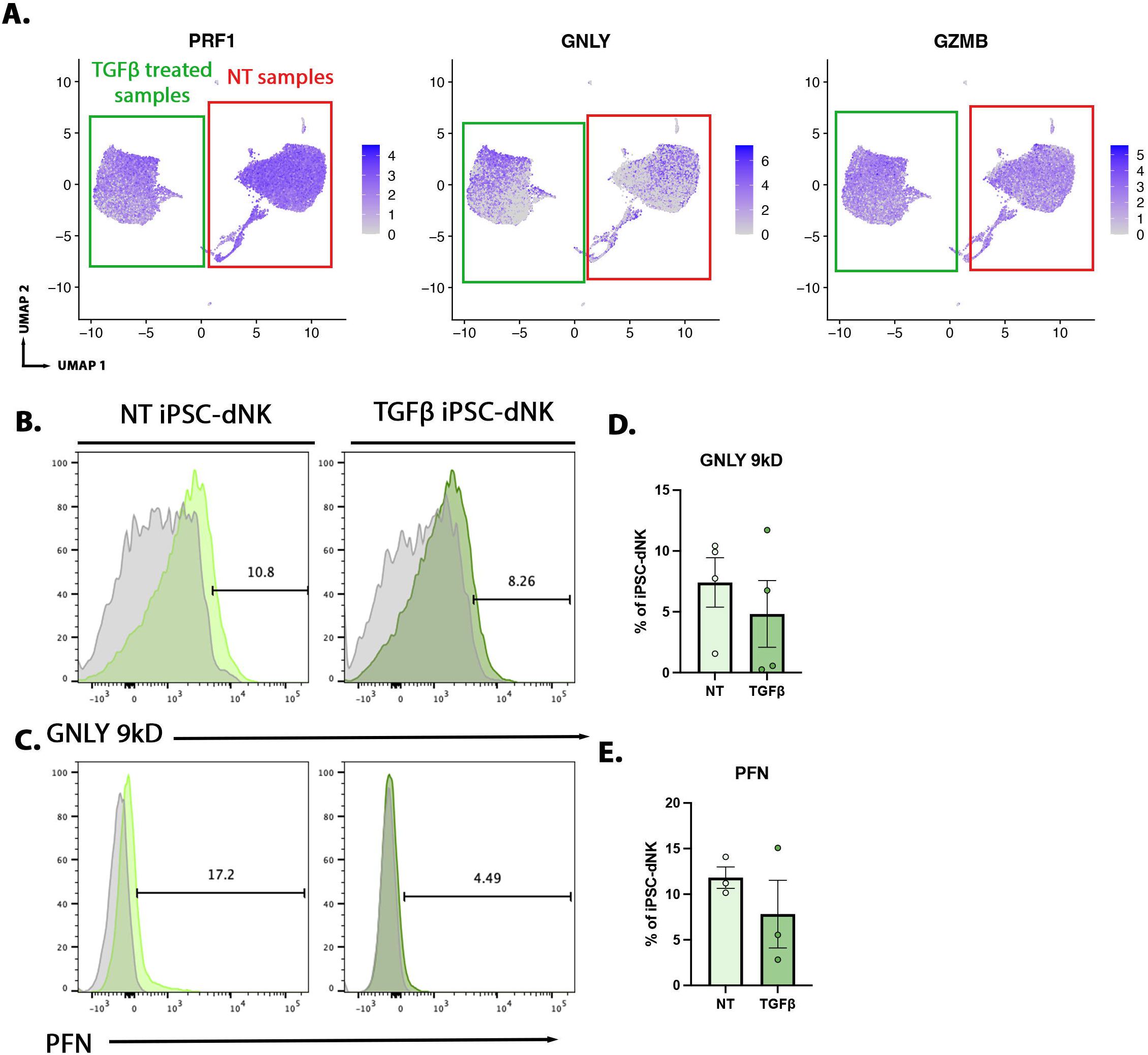

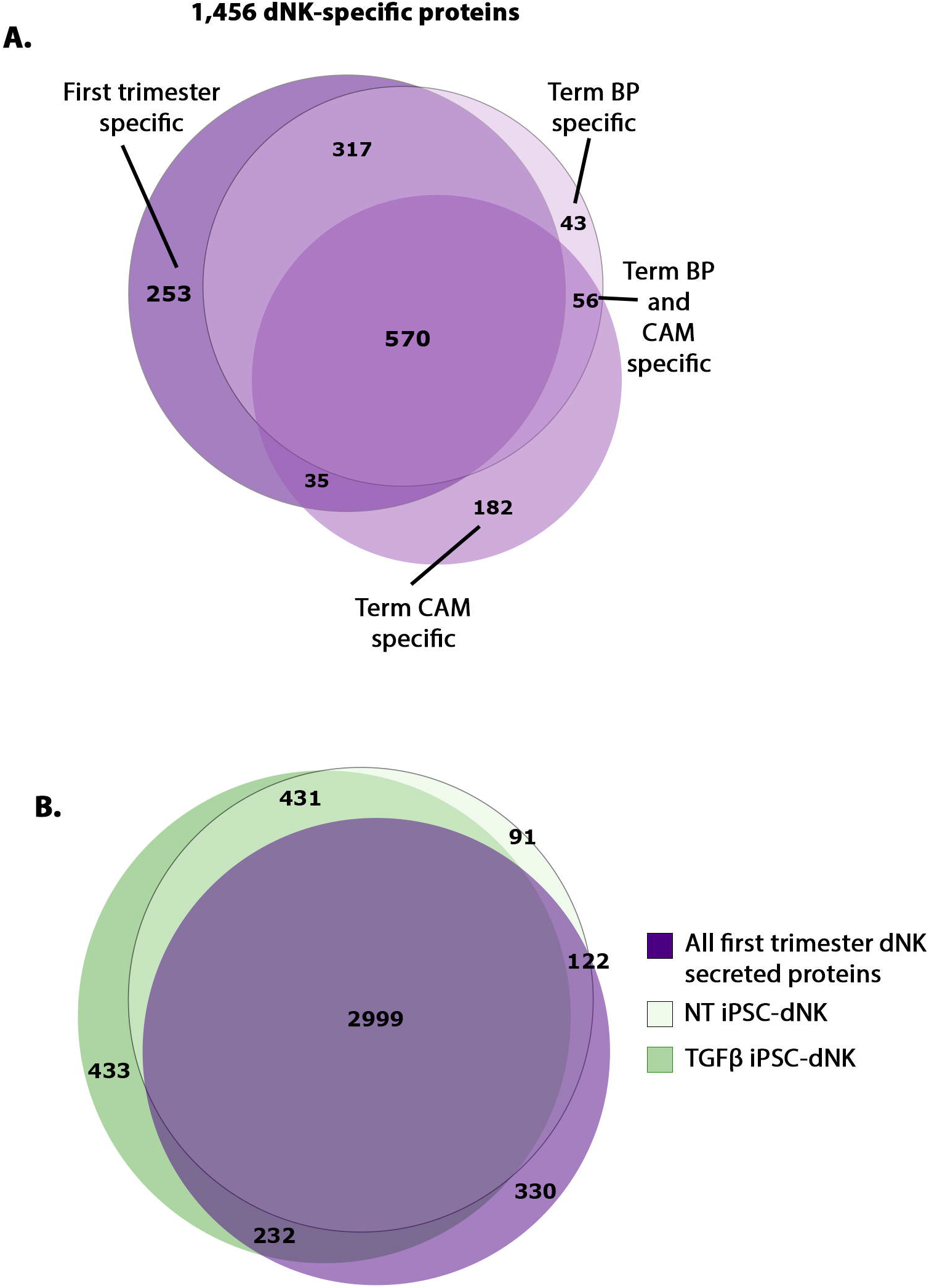

