## Supplementary Figure Legends for "Derivation of functional early gestation decidual natural killer cell subtypes from induced pluripotent stem cells"

**Running head of title: induced pluripotent stem cell-derived decidual natural killer cells.**

**Supplementary Information**

**Author list:** Virginia Chu Cheung[1,2,3], Jennifer Jaimez[1,2,3], Carly DaCosta[1,2,3], Harneet Arora[1,2,3], Christine Caron[1,3], Jaroslav Slamecka[1,2,3], Manuel Fierro[2,5], Morgan Meads[1,2,3], Kathleen Fisch[3,4], Robert E. Morey[1,2,3], Luisjesus S. Cruz[2,5], Devika Pant[1,2,3], Dan S. Kaufman[2,5], Mariko Horii[1,2,3], Jack D. Bui[1,3,*], Mana M. Parast[1,2,3,*]

[1] Department of Pathology, School of Medicine, University of California San Diego, La Jolla, CA,92093, USA.

[2]Sanford Consortium for Regenerative Medicine, University of California San Diego, La Jolla, CA,92093, USA.

[3]Center for Perinatal Discovery, University of California San Diego, La Jolla, CA, 92093, USA.

[4]Department of Obstetrics, Gynecology, and Reproductive Sciences, School of Medicine, University of California San Diego, La Jolla, CA, 92093, USA.

[5]Department of Medicine, School of Medicine, University of California San Diego, La Jolla, CA, 92093, USA.

**Disclaimers**: V.C.C., J.J., C.D., H.A., C.C., J.S., M.F., M.M., K.F., R.E.M., L.S.C., D.P., M.H., J.D.B, and M.M.P declare no competing interests. D.S.K. is a co-founder and advisor to Shoreline Biosciences and has an equity interest in the company. D.S.K. also consults Therabest and RedC Bio for which he receives income and/or equity. Studies in this work are not related to the work of those companies. The terms of these arrangements have been reviewed and approved by the University of California, San Diego, in accordance with its conflict-of-interest policies.

**Supplementary Figure Legends:**

**Supplementary Figure 1. scRNA-seq analyses of dNK subtypes in term placentas – related to Figure 1. A)** UMAP showing single cell RNA-seq (scRNA-seq) reference data containing peripheral blood, first trimester placenta, and first trimester decidua from Vento-Tormo, 2018[22] as annotated by the authors. **B)** FeaturePlot identifying NK cell populations in Vento-Tormo, 2018[22] dataset based on expression of NK cell markers *NKG7, PTPRC*, *NCAM1*. **C)** Top DEGs distinguishing first trimester dNK and PB-NK subtypes. **D)** UMAP showing term placental scRNA-seq data, containing cells isolated from placental villous tissue, basal plate, and chorioamniotic membranes from Pique-Regi, 2019[49], manually annotated using DEG information from the original publication. **E)** UMAP of Pique-Regi dataset identifying cells by tissue of origin: basal plate (BP), chorioamniotic membranes (CAM), and placental villous (PV) regions of term placenta. **F)** FeaturePlot identifying NK cell populations in Pique-Regi dataset based on expression of NK cell markers *NKG7, PTPRC*, *NCAM1.* **G)** BP and CAM NK cells extracted from entire Pique-Regi dataset and UMAP recalculated for clarity. **H)** FeaturePlots of dNK and PB-NK subtype marker genes in term BP and CAM NK cells, using specific marker genes identified in **(C)**. **I)** Bar graphs showing increased proportion of dNK2-specific $ZNF683^{+}$ cells in CAM-NK cells and increased proportion of dNK3-specific $CCL5^{+}$ cells in BP-NK cells. **J)** Pie charts showing dNK cell subtype composition (dNK1, dNK2, dNK3, and dNKp) in Pique-Regi, 2019[49] dataset by placental compartment and labor group – term in labor, TIL; term no labor, TNL; preterm labor, PTL. Of note $\leq$0.1% of all term dNK mapped to dNKp.

**Supplementary Figure 2. Newly generated induced pluripotent stem cells (iPSC) from peripheral blood mononuclear cells (PBMCs)**. **A)** Schematic illustrating the reprogramming procedure. **B)** FACS plot of isolated PBMCs showing a typical forward and side scatter profile. **C)** Digital karyotyping (by SNP array) of the three newly-generated iPSC, showing no chromosomal abnormalities after reprogramming. **D)** Bright phase images showing expected iPSC clonal morphology (10x magnification). **E)** Representative FACS plots of the three newly-generated iPSC lines showing expression of pluripotency marker SSEA4.

**Supplementary Figure 3. Identification of dNK-specific cell signaling pathways to promote a dNK cell-like phenotype from iPSC- related to Figure 2. A)** UMAP showing single cell RNA-seq (scRNA-seq) reference data from Vento-Tormo, 2018.[22] Annotations for NK cell populations were consolidated to dNK (containing dNK1, dNK2, dNK3, dNK p) and PB-NK (containing $CD16^+$ PB-NK and $CD16^-$ PB-NK). **B)** Pathway analysis of DEGs generated by direct comparison of dNK and PB-NK populations from **(A)**. **C-D)** Flow cytometry-based expression of dNK signature proteins **(C)** and dNK-associated proteins which change throughout gestation **(D)** in iPSC-derived NK cells maintained in NK differentiation media (without expansion media), No treatment (NT, expansion with IL2 and IL15 only), CXCL12 (expansion with IL2, IL15, and CXCL12), or TGFβ (expansion with IL2, IL15, and TGFβ) in 5% or 21% $O_2$. Data are represented as mean +/- standard error of n=3 separate experiments with MB3144 iPSC. Ordinary one-way ANOVA with multiple comparisons using GraphPad Prism; *p<0.05, **p<0.01, ***p<0.001, ****p<0.0001.

**Supplementary Figure 4. scRNA-seq analysis of iPSC-NK cells - related to Figure 3. A)** FeaturePlots showing expression of dNK and PB-NK subtypes marker genes in iPSC-derived NK cells identified in **Supplementary Figure 1H. B)** Bar graph showing the number of cells represented by each cluster. **C)** FeaturePlots of iPSC-derived NK cells showing expression of TGFβ-activated genes *JUN* and *SMAD7*. **D)** UMAP showing cell cycle phase predictions for iPSC derived NK. **E)** FeaturePlots of iPSC-derived NK cells showing expression of genes related to cell proliferation – *MKI67* and *PCNA*. **F)** UMAP of dNK subtypes – dNK1, dNK2, dNK3, dNKp – subsetted and reclustered from the Vento-Tormo, 2018 dataset[22], used as reference data for **Figure 3H**. **G)** Prediction scores for reference mapping results shown in **Figure 3H.**

**Supplementary Figure 5. Cytolytic molecule expression in iPSC-dNK cells - related to Figures 4 and 5. A)** FeaturePlot showing gene expression of cytolytic molecules *PRF1*, *GNLY*, and *GZMB* in NT and TGFβ iPSC-dNK cells. Representative FACS plots of cytolytic proteins GNLY 9kD **(B)** and PFN **(C)** in NT and TGFβ iPSC-dNK cells as measured by flow cytometry. **D)** Quantification of GNLY 9kD from NT and TGFβ iPSC-dNK (n=4 independent differentiations of MB3140 and MB3144 iPSC lines). **E)** Quantification of PFN from NT and TGFβ iPSC-dNK cells (n=3 independent differentiations of MB3140 and MB3144 iPSC lines). Statistical testing in (**D**) and (**E**) of NT vs. TGFβ iPSC-dNK were compared by paired student's t-test using GraphPad Prism.

**Supplementary Figure 6. Comparison of whole secretomes of primary first trimester and term dNK cells. – Related to Figure 4 A)** Venn Diagram representation of dNK cell-specific proteins (total of 1,456) identified in **Figure 4B** by sample group - first trimester, term BP, or term CAM dNK cells. Proteins found in conditioned media from first trimester, but not in term BP or term CAM, dNK cells were identified as first trimester-specific (253). Term BP-specific proteins (43) were not present in first trimester or term CAM dNK cell conditioned media. Term CAM-specific proteins (182) were not present in first trimester or term BP conditioned media. **B)** Venn Diagram representation of overlap between all 3,683 proteins secreted by first trimester dNK and all 3,643 NT iPSC-dNK and 4,095 TGFβ iPSC-dNK secreted proteins.

<u>**Supplementary File Information:**</u>

**Supplementary File 1 related to Table 1 and Supplementary Figure 1.** Complete list of marker genes distinguishing dNK cell subtypes (dNK1, dNK2, dNK3, dNKp) and PB-NK cell subtypes (PB-NK $CD16^+$, PB-NK $CD16^-$).

**Supplementary File 2 related to Figure 3.** Marker genes distinguishing iPSC-dNK cells clusters.

**Supplementary File 3 related to Figure 3.** Pathway analysis of NT clade, TGFβ clade, and proliferative clade from iPSC-dNK cell marker genes using MSigDB_Hallmark_2020 library.
