## Supplementary Materials and Methods for "Derivation of functional early gestation decidual natural killer cell subtypes from induced pluripotent stem cells"

**Running head of title**: **induced pluripotent stem cell-derived decidual natural killer cells.**

### Supplementary Information

**Author list:** Virginia Chu Cheung[1,2,3], Jennifer Jaimez[1,2,3], Carly DaCosta[1,2,3], Harneet Arora[1,2,3], Christine Caron[1,3], Jaroslav Slamecka[1,2,3], Manuel Fierro[2,5], Morgan Meads[1,2,3], Kathleen Fisch[3,4], Robert E. Morey[1,2,3], Luisjesus S. Cruz[2,5], Devika Pant[1,2,3], Dan S. Kaufman[2,5], Mariko Horii[1,2,3], Jack D. Bui[1,3,*], Mana M. Parast[1,2,3,*]

[1] Department of Pathology, School of Medicine, University of California San Diego, La Jolla, CA,92093, USA.

[2]Sanford Consortium for Regenerative Medicine, University of California San Diego, La Jolla, CA,92093, USA.

[3]Center for Perinatal Discovery, University of California San Diego, La Jolla, CA, 92093, USA.

[4]Department of Obstetrics, Gynecology, and Reproductive Sciences, School of Medicine, University of California San Diego, La Jolla, CA, 92093, USA.

[5]Department of Medicine, School of Medicine, University of California San Diego, La Jolla, CA, 92093, USA.

**Disclaimers**: V.C.C., J.J., C.D., H.A., C.C., J.S., M.F., M.M., K.F., R.E.M., L.S.C., D.P., M.H., J.D.B, and M.M.P declare no competing interests. D.S.K. is a co-founder and advisor to Shoreline Biosciences and has an equity interest in the company. D.S.K. also consults Therabest and RedC Bio for which he receives income and/or equity. Studies in this work are not related to the work of those companies. The terms of these arrangements have been reviewed and approved by the University of California, San Diego, in accordance with its conflict-of-interest policies.

### Supplementary Materials and Methods

#### Patient recruitment and tissue collection

Human decidual and blood tissue samples for this study were collected under UC San Diego Institutional Review Board-approved protocols (IRB #181917 and 172111); all patients gave informed consent for collection and use of these tissues. First trimester samples were collected from pregnant individuals, undergoing elective termination of pregnancy, providing written informed consent. First trimester decidual tissues, were collected from patients with gestational ages ranging from 5 to 9 weeks (n=4). Term basal plate and chorioamniotic tissues (n=4) were collected from non-laboring patients undergoing elective c-section 39w-39w5d gestational age. Gestational age was determined based on crown-rump length, measured by first trimester ultrasound, and was stated in weeks/days from the first day of the last menstrual period. "Normal" sample was defined as a singleton pregnancy without any detectable fetal abnormalities on ultrasound; for term samples, the definition also included absence of chronic maternal conditions (e.g. autoimmune disease, chronic hypertension) or pregnancy complications (e.g. gestational diabetes, preeclampsia).

#### Primary dNK preparation

First trimester decidual fragments were finely minced and digested in RPMI (Corning) containing 1mg/mL of Collagenase IA (Sigma-Aldrich) and 0.2mg/mL of DNase I (Roche) for 15-30min in a shaking incubator at 37°C at 200rpm. Digested tissue was washed using RPMI with 10% FBS (Cell Culture Collective, Inc.) and 0.2mg/mL of DNase I and filtered through 100um, 70um, then 40um filters. Flow through was collected and pelleted by centrifugation at 400g for 5min. The pelleted cells were resuspended in a 25% (final concentration 1.029 g/mL Percoll solution and layered on top of a 1.082 g/mL and 1.053 g/mL Percoll layers. The gradient was topped with 1x HBSS. The gradient was centrifuged at 800g for 30min with both the acceleration and deceleration set to 0. After centrifugation, the lymphocyte layer was collected from the 1.082/1.053 g/mL interface. Cells were washed and either stained and analyzed by flow cytometry or purified further by NK Cell MACS negative selection (Miltenyi Biotec) for functional analyses.

Term placental basal plate was dissected from a 2-3mm depth of the maternal surface and minced to isolate dNK. The maternal surface of the chorioaminotic membranes was scraped to isolate dNK. Both tissues were digested in RPMI containing 1mg/mL of Collagenase IA and 0.2mg/mL of DNase I for 30min in a shaking incubator at 37°C at 200rpm. Digested tissue was washed using RPMI with 10% FBS and 0.2mg/mL of DNase I and passed through an initial metal sieve to remove undigested tissues. The flow-through was filtered through 100µm, 70µm, then 40µm filters. Flow through was collected and pelleted by centrifugation at 400g for 5min. The pelleted cells were resuspended in a 25% (final concentration 1.029 g/mL) Percoll solution and layered on top of 1.082 g/mL and 1.053 g/mL Percoll layers. The gradient was topped with 1x HBSS. The gradient was centrifuged at 800g for 30min with both the acceleration and deceleration set to 0. After centrifugation, the lymphocyte layer was collected from the 1.082/1.053 g/mL interface. Cells were washed and either stained and analyzed by flow cytometry or purified further by FACS using an Aria II sorter ($CD3^{-}CD19^{-}CD15^{-}CD45^{+}CD56^{+}$) for functional analyses.

**<u>Peripheral Blood Mononuclear Cell sample collection</u>**

Human blood samples were collected from non-pregnant (proteomics or pregnant (XX) individuals in the third trimester. Whole blood collected in K2 EDTA tubes and centrifuged at 1200 x g for 15 min with deceleration set to 1 at room temperature. The buffy coat layer containing peripheral blood mononuclear cells (PBMCs) is collected and washed in 1x PBS (Corning). Red blood cell contamination was removed by ACK cell lysis buffer (Gibco) for 7min at 37°C. PBMCs were washed again and frozen in aliquots of 1-2 x $10^6$ cells per cryovial in 90% Knockout Serum Replacement (Gibco) and DMSO (Sigma-Aldrich).

**Generation of iPSC lines**

Reprogramming was done using CytoTune -iPSC 2.0 Sendai Reprogramming Kit (Thermo Fisher Sciencetific cat # A16517) following the manufacturer's protocol. In brief, reprogramming vectors were transduced into PBMCs collected from pregnant (XX) individuals after culture in StemPro34 media (Thermo Fisher Scientific, cat# 10639011) containing SCF, IL6, IL3, FLT3. Post-transduction day 5–7, PBMCs were replated onto Geltrex (Gibco cat #A14133-02)-coated dishes with mTeSR Plus medium (StemCell Technologies, cat# 100-0276). iPSC colonies appeared around 2 weeks post-transduction, and were subsequently picked for subcultures and passaged up to 10 times to dilute Sendai virus out from the culture. All the cells used in this study passed quality control assays, including confirmation of pluripotency (based on morphology and flow cytometry for SSEA4) (**Supplementary Figure 2D,E**). Cells were collected from 1 70-80% confluent well of a 6 well plate for genomic DNA isolation (Qiagen #69506) for SNP genotyping at passage 10. We used SNP genotyping (Infinium Core-24, Illumina) to confirm that 3 different individual iPSC lines (MB3140, MB3144, and LD2809) were used in this study and that they had a normal karyotype (**Supplementary Figure 2C**). Finally, we used PCR to confirm the absence of residual expression of exogenous reprogramming factors (data not shown) prior to differentation. Cells were frozen at or before passage 10 in a cryovial containing Freezing Media (90% Knockout Serum Replacement (KSR, Thermo Fisher Scientific cat#10-828-028) and 10% DMSO (Sigma Aldrich cat # D2650-100ML)) and stored in liquid nitrogen. To thaw, cryovials were warmed in a 37C water bath and quickly resuspended in mTeSR Plus complete. Cells were removed from Freezing Media by centrifugation at 300xg for 5min. Pelleted cells were resuspended in mTeSR Plus complete and plated onto a Geltrex coated tissue culture plate. All cells were cultured using sterile technique in BSL2 tissue culture hood and were routinely tested for mycoplasma using MycoAlert Detection Kit (Lonza, cat# LT07-318) and confirmed negative for all established lines.

**hPSC culture and differentiation to dNK-like cells**:

The human pluripotent stem cell (hPSC) aspect of the research was performed under a protocol approved by the University of California, San Diego Institutional Review Board and Embryonic Stem Cell Research Oversight Committee (#171648). Before differentiation, hESC WA09/H9 and 3 iPSC lines (MB3140, MB3144, and LD2809, established as a part of the biorepository of the Center for Perinatal Discovery) were used for differentiation with specific sample number information provided for each result reported. Cell lines were handled separately by a single individual at any given time. Individual cell lines were maintained in separate tissue culture vessels with the top and bottom of the plates labelled. hPSC lines were maintained in mTeSR Plus on Geltrex-coated plates. hPSC were passaged at least 2 passages from time of thaw prior to differentiation using 0.5mM EDTA (Life Technologies cat # 15575-08) in PBS at plated at appropriate split ratios (1:3-1:6). iPSC at passage 20 or earlier and hESC at passage 50 or earlier were used for differentiation. For differentiation into dNK, we adapted a previously-published protocol for differentiation into NK (Zhu, 2019). In brief, hPSC with good purity (100% SSEA-4+) were dissociated to single cell using TryPLE and resuspended at a density of $1x10^5$ cells/mL in APEL2 (STEMCELL Technologies) containing 40ng/mL of SCF, 20ng/mL of BMP4, 20ng/mL VEGF, and 10uM of Y27632. 100,000 cells were plated into each well of a U-bottom 96-well plate and centrifuged at 300 g for 5 minutes. Cells were cultured for 6 days to allow EBs to form. At day 6, EBs were evaluated for CD34 expression by flow cytometry and collected for NK differentiation. 20-30 EBs were seeded into one well of a 6-well plate in NK Differentiation Media (DMEM/F12, 15% heat-inactivated human AB serum (Valley Biomedical: HP1022HI), 1x Glutamax. (Life Technologies # 35050061), 27.5uM β-mercaptoethanol (Life Technologies # 21985-023), 5 ng/mL sodium selenite (Sigma Aldrich: S5261-10G), 50 μM ethanolamine (MP Biomedicals:

0219384590), 20μg/mL L-ascorbic acid 2-phosphate (Sigma Aldrich: A8960-5G) containing 5ng/mL IL-3 (PeproTech: 200-03), 20ng/mL SCF (PeproTech: 300-07), 20ng/mL IL-7 (PeproTech: 200-07), 10ng/mL IL-15 (PeproTech:200-15), 10ng/mL FLT3 ligand (PeproTech: 300-19). Media was changed every 3-5 days after initial seeding with NK Differentiation media containing SCF, IL-7, IL-15, and FLT3 ligand for a total of 28-42 days. After 28 days in the NK Differentiation phase, we confirmed the commitment to NK cells by measuring expression of CD45, CD56, and CD16. We expected 80-100% of the cells to be $CD45^+$, 30-70% to be $CD56^+$, and all the cells to be $CD16^-$. To expand the cells, we collected the cells from the NK Differentiation media by centrifugation at 600g for 5min. We removed the NK Differentiation media and resuspended the cells in CTS Xpander media (Thermo Scientific: A5019001) according to the manufacturer's recommendation of 500,000 cells per 1mL for an additional 2 or 3 weeks (specific length of time noted for each experiment) in the presence of 500units/mL of IL-2 (PeproTech: 200-02) and 10ng/mL of IL-15 at atmospheric (21%) $O_2$ unless otherwise noted. 10ng/mL (PeproTech: 100-21) TGFβ1 or 100ng/mL CXCL12 (PeproTech: 300-28A) were included in some experiments as detailed in the results. Cultures maintained at 5% $O_2$ were incubated and maintained in an XVIVO Model X3 Hypoxia Workstation (Biospherix, Redfield, NY). Cells were visualized and images acquired using an EVOS XL Core and EB images were acquired using an Olympus dissection microscope (Model SZX2-ILLD). After 3 weeks of treatment with or without TGFβ, iPSC-NK were stimulated with PMA/I.

**PMA/I stimulation:**

Cells were plated in 5% Human AB serum in RPMI and low dose (2.5ng/mL IL-15) overnight (~16hr) with 25ng/mL PMA and 1ug/mL ionomycin. Four hours prior to collection, Brefeldin A to was added to all wells as well as 250ng/mL of either CD107a-BV510 or MsIgG-BV510 isotype antibodies. Two hours prior to collection, PMA (25ng/mL) and Ionomycin (1μg/mL) or media only (as an unstimulated control) was added. After 2 hours, cells were collected for intracellular staining.

### Flow Cytometry

#### Cell Surface staining

Single cell suspensions were collected in FACS Buffer (10% FBS, 5mM EDTA in PBS). Proteins were stained at 4°C for 20 min protected from light. Stained cells were washed, fixed in 2% PFA and stored in FACS Buffer prior to analysis. A complete list of antibodies can be found in **Supplementary Table 4**. Stained samples were analyzed using BD FACS Canto.

#### Intracellular staining

Collected cells were blocked for 15 min at room temperature (RT) protected from light using 10% Human AB Serum in PBS. After blocking, cells were washed and Zombie fixable live/dead stain was applied at a 1:1000 dilution in PBS. After 20min at RT protected from light, cells were washed and cell surface proteins were stained for in BD Brilliant Stain Buffer (BD Biosciences cat #563794) at 4°C for 20 min protected from light. Cells were washed and fixed and permeabilized using 1x BD Perm/Wash buffer. Intracellular cytokines were stained for in 1x BD Perm/Wash buffer with a 30 min incubation at 4°C protected from light. Stained cells were washed and stored in FACS Buffer (10% FBS, 5mM EDTA in PBS) prior to analysis. A complete list of antibodies can be found in **Supplementary Table 4**. Stained samples were analyzed using BD Fortessa X20.

### Aptamer-based proteomic profiling

200k purified primary cells or iPSC-NK cells expanded with or without TGF$\beta$, were plated in 500uL of 5% Human AB serum in RPMI and low dose (2.5ng/mL) IL-15 in a 24-well non-TC treated plate overnight (~20hr). Cells and media were collected and separated by centrifugation at 600g for 5min. Media was stored in the -80°C until analysis. Proteomic profiling was performed using the

SomaScan Assay V4.1 (SomaLogic, Boulder, CO, USA) 7K panel, a high-throughput proteomics platform that employs SOMAmer (Slow Off-rate Modified Aptamer) Reagents to quantify the relative concentrations of proteins. Assays were conducted at SomaLogic's headquarters (Boulder, CO, USA) under the Standard BioTools Quality System (QS). The laboratory operates in compliance with Clinical Laboratory Improvement Amendments (CLIA) standards for laboratory-developed tests and is accredited to ISO 15189 standards for medical laboratories. Quality control Relative protein abundances are expressed in relative fluorescence units (RFU), fully normalized following a standard procedure to account for nuisance effects in hybridization, sample volume and plate biases, and $log_{10}$-transformed after normalization. Proteins with RFUs above media only control were selected for analysis. Individual proteins were compared between all seven sample groups using Kuskal-Wallis test followed by Dunn's test for multiple comparisons. Scaled Venn Diagrams were calculated using DeepVenn.[68]

**<u>Kill Assay:</u>**

**Effector cells:**

<u>Primary dNK:</u>

Percoll purified lymphocytes from first trimester decidua, term BP, and term CAM were cryopreserved in 90% KSR/10% DMSO and stored in liquid nitrogen. Percoll-purified lymphocytes were thawed and first trimester dNK were purified by NK Cell MACS negative selection (Miltenyi Biotec). Term dNK were purified by FACS (Aria Fusion Sorter) using 7-AAD$^-$CD3$^-$CD15$^-$CD19$^-$ CD56$^{bright}$. After purification by either MACS or FACS, primary dNK were resuspended in media (5% Human AB Serum in RPMI with 2.5ng/mL IL-15) and recovered at 37°C in 5% $CO_2$ for 6hrs prior to coculture with target cells. >90% viability was confirmed by trypan blue exclusion before coculture and cells were counted used the Countess FL3. Cells were counted and >90% viability was confirmed by trypan blue exclusion using the Countess FL3.

<u>iPSC-dNK</u>

NT and TGFβ-treated iPSC-dNK were generated as described above. Cells were counted and >90% viability was confirmed by trypan blue exclusion using the Countess FL3.

PB-NK

Peripheral blood mononuclear cells (PBMC) were isolated through density gradient centrifugation from an apheresis product (obtained from anonymized donors from the San Diego Blood Bank through an IRB exempt protocol) and NK cells were sorted using EasySep Human NK Cell Enrichment Kit (StemCell Technologies, cat# 17955) following manufacture protocol from 3 different donors. Peripheral blood NK cells were cultured in RPMI 1640 (Invitrogen), 10% heat-inactivated human AB Serum (BioIVT, cat# HUMANABSRMP-HI-1), 1X Glutamax (Thermo Scientific cat# 35050061), 1% PenStrep (Fisher Scientific, cat# 15140122) and supplemented with 100 IU/mL of hIL-2 and 10ng/mL hIL-15 every three days or media change. NK cells were co-cultured once a week with irradiated artificial antigen-presenting cells for expansion. Cells were counted and >90% viability was confirmed by trypan blue exclusion using the Countess FL3.

NK-92

NK92 cell line was purchased from ATCC and cultured in media containing 12% Horse Serum (Life Technologies, cat# 16050122), 12% FBS (Millipore Sigma, cat# F4135), 1x Glutamax (Thermo Scientific cat# 35050061), 200uM Myoinositol (Sigma-Aldrich, cat# I7508), 20uM Folic Acid (Sigma-Aldrich, cat# F8758), 100uM beta-mercaptoethanol (Life Technologies, cat# 21985023) in alpha-MEM without ribo/deoxyribonucleosides (Gibco, cat# 12561056) supplemented with 100 IU/mL of hIL-2 and 10ng/mL hIL-15. Cells were counted and >90% viability was confirmed by trypan blue exclusion using the Countess FL3.

**Target cells**

JEG3 cells were purchased from ATCC and were cultured in media containing 10%FBS (Omega scientific cat# FB-02 in DMEM, high glucose Gibco (Thermo Fisher cat#11965092). Cells at 70-80% confluence were dissociated using 0.05% Trypsin with EDTA (Corning cat # 25-051-CI).

Cells were counted and >90% viability was confirmed by trypan blue exclusion using the Countess FL3.

**Assay**

All effector and Target cells were cocultured in 200uL of media (5% Human AB Serum in RPMI with 2.5ng/mL IL-15) at 1:1, 2:1, and 5:1 E:T ratios for 12hrs at 37°C in 5% $CO_2$. Target only controls were included in every time the assay was performed. After 12 hours, cells were washed with 2mL of 1x PBS and stained for apoptotic cells using the Dead Cell Apoptosis Kit with Annexin V Alexa Flour 488 & Propidium Iodide (Thermo Scientific Cat #V13241). Effector and target cells were distinguished using EGFR and CD45 antibodies. (**Supplementary Table 4**).

### Single cell RNA-sequencing reanalysis

First Trimester, Peripheral Blood – Vento-Tormo, 2019

The Seurat object containing data from all the published tissues (first trimester decidua, first trimester placenta, and peripheral blood) processed by both 10x 3' v2 and Smart-seq2 with all associated metadata was obtained from cellxgene (https://cellxgene.cziscience.com/collections/a9254216-6cd8-4186-b32c-349363777584). Counts were extracted, log normalized and scaled using 10,000 variable features identified by FindVariableFeatures. PCA and UMAP were calculated using Seurat v5.0.3[56]. Instead of performing clustering analysis, provided "author_cell_type" annotations were used with the exception of T cells. T cell annotations were updated with "tissue" information to result in "Tcell_decidua", "Tcell_placenta", and "Tcell_blood" to better reflect the published clustering.

All NK cells (dNK1, dNK2, dNK3, dNKp, NK CD16+, and NK CD16-) were extracted from this dataset and a new UMAP visualization was calculated to better visualize the relevant cells as a reference for Figure 1A. NK subtype-specific markers were identified using the FindAllMarkers function of Seurat with the parameter 'method= wilcox, only.pos = TRUE, logfc.threshold = 0.1'.

Marker genes were further filtered to reflect genes expressed by greater than 30% of cells in a given NK subtype and less than 50% of all other NK cells (pct.1 >0.3, pct.2<0.5).

In select analyses, we updated the annotations to reflect broad clustering of dNK cells (dNK1, dNK2, dNK3, and dNKp together) and PB-NK cells (NK $CD16^{+}$ and NK $CD16^{-}$). dNK cell-specific pathways were identified by inputting marker genes calculated by FindMarkers function of Seurat with the parameter ‘method= wilcox, only.pos = TRUE, logfc.threshold = 0.1’comparing dNK cell (pct.1) and PB-NK cell (pct.2). dNK marker genes were further filtered to reflect genes expressed by greater than 80% of dNK (pct.1>0.8) and inputted to Enrichr and queried against the MSigDB_Hallmark_2020 library.

Term – Pique-Regi, 2019

Raw data associated with placental villous (PV), basal plate (BP), and chorioamniotic membrane (CAM) tissues from Pique-Regi, 2019 (Version 1) were downloaded with permission from **dbGaP Study Accession:** phs001886.v3.p1. The preliminary filtered data generated from Cell Ranger (Cell ranger v7.1.0) were used for downstream filtering and Seurat v5.0.3[65] was used for analysis. Each sample was annotated with associated meta data – specifically, placenta location (BP, CAM, PV), and labor group (TIL, TNL, PTL). First, we filtered out cells expressing less than 200 genes and selected genes expressed by more than 10 cells. We further filtered the dataset for cells expressing less than 20% mitochondrial genes, greater than 5% ribosomal genes, and less than 0.05% hemoglobin genes. The filtered dataset was normalized (normalization.method = “LogNormalized”, scale.factor = 10,000) and scaled. PCA and UMAP were calculated using using 2,000 variable features identified by FindVariableFeatures (Seurat v5.0.3). Clustering was performed using FindNeighbors (dims = 1:30) and FindClusters (resolution = 0.5). The published UMAP and DEG information were used to guide cluster resolution selection and manual annotation. BP and CAM samples, which include the maternal compartments of the placenta containing dNK cells, were subsetted for reference mapping analyses. After predicted identities were calculated for each cell, associated meta data such as decidual location (BP vs. CAM) and labor group (TIL, TNL, PTL) were used for select analyses.

**scRNA-seq library generation and analysis of iPSC-NK**

Cells from two iPSC lines were differentiated to iPSC-NK cells and, after 2 weeks of expansion with or without TGFβ, were run on the 10X Genomics platform with the Chromium Next GEM Single Cell 3' v3 kit. Some cells were reserved to verify expression of CD45, CD56, CD16, CD9, and CD103 prior to library preparation. Samples and libraries were prepared following the manufacturer's protocol to target 10,000 cells per reaction. Libraries were pooled and sequenced using the Illumina NovaSeq 6000 sequencer.

The preliminary filtered data generated from Cell Ranger (Cell ranger v7.2.0) was used for downstream filtering and Seurat v5.0.3[65] was used for analysis. Each sample was annotated with associated meta data – specifically, treatment group (TGFβ or NT), and cell line (MB3144 and MB3140). First, we filtered out cells expressing less than 200 genes and selected genes expressed by more than 10 cells. We further filtered the dataset for cells expressing less than 20% mitochondrial genes, between 5% and 25% ribosomal genes, and less than 0.05% hemoglobin genes. Doublets were identified and removed using DoubletFinder.[66] The filtered dataset was normalized (normalization.method = "LogNormalize") and scaled using variable features identified by FindVariableFeatures. PCA and UMAP were calculated using Seurat v5.0.3. Due to the long duration of TGFβ treatment and the expression pattern of dNK cell marker proteins CD9 and CD103, we determined that iPSC-NK cells derived with and without TGFβ treatment were distinct populations. To analyze the difference between treatment groups and avoid differences driven by cell line, we integrated our data using CCA integration with "cell line" as a covariate. Clustering after integration was performed using FindNeighbors (dims = 1:30) and

FindClusters (resolution = 0.3). Clustree[67] and Seurat's BuildClusterTree was used to guide the selection of a cluster resolution.

Cluster-specific markers were identified by using the FindAllMarkers function of Seurat with the parameter 'method= wilcox, only.pos = TRUE, logfc.threshold = 0.1'. Marker genes were further filtered to reflect genes expressed by greater than 50% of cells in a given cluster and less than 50% of all other cells (pct.1 >0.5, pct.2<0.5).

Pathway analysis was performed by inputting cluster marker genes grouped by clade to Enrichr using the MSigDB_Hallmark_2020 and Descartes_Cell_Types_and_Tissue_2021 libraries.

**Statistical Tests**

All statistical analysis of flow cytometry data was performed using GraphPad Prism 10.

All comparisons of flow cytometry data between three groups were tested using Ordinary one-way ANOVA with multiple comparisons. Comparisons between NT and TGF$\beta$ groups were made using Paired student's t-test.

Statistical analysis of Annexin $V^+$ expression between seven effector groups (column factor) and 3 E:T ratios (row factor) were performed using Two-way ANOVA with Sidak's test for multiple comparisons using GraphPad Prism 10.

All cell culture experiments were performed at least three independent replicates with detailed sample information provided in the figure legend for each experiment. Aptamer-based proteomic profiling data was determined to have a not-normal distribution using the Anderson-Darling normality test and therefore comparisons between groups were made using Kuskal-Wallis followed by Dunn's test for multiple comparisons using R.
