## Supplementary Tables for "Derivation of functional early gestation decidual natural killer cell subtypes from induced pluripotent stem cells"

**Running head of title: induced pluripotent stem cell-derived decidual natural killer cells.**

### Supplementary Information


**Author list:** Virginia Chu Cheung[1,2,3], Jennifer Jaimez[1,2,3], Carly DaCosta[1,2,3], Harneet Arora[1,2,3], Christine Caron[1,3], Jaroslav Slamecka[1,2,3], Manuel Fierro[2,5], Morgan Meads[1,2,3], Kathleen Fisch[3,4], Robert E. Morey[1,2,3], Luisjesus S. Cruz[2,5], Devika Pant[1,2,3], Dan S. Kaufman[2,5], Mariko Horii[1,2,3], Jack D. Bui[1,3,*], Mana M. Parast[1,2,3,*]

[1] Department of Pathology, School of Medicine, University of California San Diego, La Jolla, CA,92093, USA.

[2]Sanford Consortium for Regenerative Medicine, University of California San Diego, La Jolla, CA,92093, USA.

[3]Center for Perinatal Discovery, University of California San Diego, La Jolla, CA, 92093, USA.

[4]Department of Obstetrics, Gynecology, and Reproductive Sciences, School of Medicine, University of California San Diego, La Jolla, CA, 92093, USA.

[5]Department of Medicine, School of Medicine, University of California San Diego, La Jolla, CA, 92093, USA.


**Disclaimers**: V.C.C., J.J., C.D., H.A., C.C., J.S., M.F., M.M., K.F., R.E.M., L.S.C., D.P., M.H., J.D.B, and M.M.P declare no competing interests. D.S.K. is a co-founder and advisor to Shoreline Biosciences and has an equity interest in the company. D.S.K. also consults Therabest and RedC Bio for which he receives income and/or equity. Studies in this work are not related to the work of those companies. The terms of these arrangements have been reviewed and approved by the University of California, San Diego, in accordance with its conflict-of-interest policies.

**Supplementary Table 1. Pathway analysis of NT clade using Descartes_Cell_Types_and_Tissue_2021 library**

| Term | # of Enriched Genes | # Term Genes | Adjusted P-value |
|---|---|---|---|
| Microglia in Cerebellum | 122 | 606 | 3.40E-14 |
| Lymphoid cells in Intestine | 48 | 156 | 3.23E-12 |
| Microglia in Eye | 56 | 240 | 4.71E-09 |
| Lymphoid cells in Placenta | 39 | 153 | 1.67E-07 |
| Lymphoid cells in Adrenal | 40 | 177 | 3.14E-06 |
| Lymphoid cells in Stomach | 23 | 85 | 4.33E-05 |
| Microglia in Cerebrum | 75 | 465 | 4.33E-05 |
| Lymphoid cells in Lung | 33 | 150 | 4.51E-05 |
| Lymphoid cells in Heart | 24 | 110 | 9.94E-04 |
| Lymphoid cells in Pancreas | 29 | 152 | 0.00217016 |

**Supplementary Table 2. Pathway analysis of TGFβ clade using Descartes_Cell_Types_and_Tissue_2021 library.**

| Term | # of Enriched Genes | # Term Genes | Adjusted P-value |
|---|---|---|---|
| Lymphoid cells in Intestine | 22 | 156 | 2.21E-06 |
| Lymphoid cells in Placenta | 20 | 153 | 1.65E-05 |
| Lymphoid cells in Adrenal | 17 | 177 | 0.00445743 |
| Lymphoid cells in Lung | 13 | 150 | 0.04792624 |
| Microglia in Eye | 17 | 240 | 0.06963168 |
| Lymphoid cells in Heart | 10 | 110 | 0.06963168 |
| Lymphoid cells in Spleen | 10 | 119 | 0.09212431 |
| Lymphoid cells in Stomach | 8 | 85 | 0.09212431 |
| Lymphoid cells in Kidney | 17 | 260 | 0.09212431 |
| Lymphoid cells in Muscle | 13 | 193 | 0.14822724 |

**Supplementary Table 3. Pathway analysis of proliferative clade using Descartes_Cell_Types_and_Tissue_2021 library**

| Term | # of Enriched Genes | # Term Genes | Adjusted P-value |
|---|---|---|---|
| Erythroblasts in Stomach | 8 | 115 | 9.47E-05 |
| Erythroblasts in Heart | 10 | 331 | 0.00414751 |
| Erythroblasts in Muscle | 5 | 191 | 0.11891546 |
| Megakaryocytes in Muscle | 4 | 288 | 0.86249075 |
| Erythroblasts in Pancreas | 2 | 140 | 0.93551307 |
| Lymphoid cells in Placenta | 2 | 153 | 0.93551307 |
| Trophoblast giant cells in Placenta | 1 | 89 | 0.99433571 |
| Lens fibre cells in Eye | 1 | 111 | 0.99433571 |
| Smooth muscle cells in Eye | 1 | 135 | 0.99433571 |
| Megakaryocytes in Lung | 1 | 191 | 0.99433571 |

**Supplementary Table 4. List of antibodies used for flow cytometry**

| Protein | Fluorochrome | Clone | Vendor | Catalog# |
|---|---|---|---|---|
| CD19 (Lineage) | PerCP-Cy5.5 | HIB19 | Biolegend | 302230 |
| CD15 (Lineage) | PerCP-Cy5.5 | W6D3 | Biolegend | 323020 |
| CD3 (Lneage) | PerCP-Cy5.5 | OKT3 | Biolegend | 317335 |
| 7-AAD | | | ThermoFisher | A1310 |
| CD45 | APC-Cy7 | HI30 | Biolegend | 304014 |
| CD45 | BV650 | HI30 | Biolegend | 304044 |
| CD45 | APC | HI30 | BD | 560973 |
| CD56 | APC | B159 | BD | 555518 |
| CD56 | PE-Cy7 | MEM-188 | Biolegend | 304628 |
| CD16 | FITC | 3G8 | Biolegend | 302006 |
| CD18 | PE-Cy7 | 1B4 | Biolegend | 373410 |
| KIR2DL1 | PE | 1127B | R&D Systems | FAB8887P-025 |
| CD9 | PE | HI9a | Biolegend | 312106 |
| CD103 | PE-Cy7 | Ber-ACT8 | Biolegend | 350212 |
| CD39 | PE | A1 | Biolegend | 328208 |
| CD69 | PE-Cy7 | FN50 | Biolegend | 310912 |
| CD49a | AlexaFlour 647 | TS2/7 | Biolegend | 328310 |
| CD34 | APC | 581 | BD | 555824 |
| CD34 | PE-Cy7 | 581 | BD | 560710 |
| SSEA4 | PE | MC-813-70 | Biolegend | 330406 |
| CD107a | BV510 | H4A3 | Biolegend | 328632 |
| IFNγ | APC/Cy7 | B27 | Biolegend | 506524 |
| TNFα | PE/Cy7 | MAb11 | Biolegend | 502930 |
| GM-CSF | Pe/dazzle™ 594 | BVD2-21C11 | Biolegend | 502318 |
| PFN | BV711 | dG9 | Biolegend | 308129 |
| GNLY 9kD | PE | DH2 | Biolegend | 348004 |
| VEGF | APC | 23410 | R&D Systems | IC2931A |
| Isotype Controls | | | | |
| Brilliant Violet 510™ Mouse IgG1, κ Isotype Ctrl Antibody | BV510 | MOPC 21 | Biolegend | 400172 |
| APC/Cyanine7 Mouse IgG1, κ Isotype Ctrl Antibody | APC-Cy7 | MOPC 21 | Biolegend | 400128 |
| PE/Cyanine7 Mouse IgG1, κ Isotype Ctrl Antibody | PE-Cy7 | MOPC 21 | Biolegend | 400126 |

| | | | | |
|---|---|---|---|---|
| PE/Dazzle™ 594 Rat IgG2a, κ Isotype Ctrl Antibody | Pe/dazzle™ 594 | RTK2758 | Biolegend | 400558 |
| Brilliant Violet 711™ Mouse IgG1, κ Isotype Ctrl Antibody | BV711 | MOPC 21 | Biolegend | 400168 |
| PE Mouse IgG1, κ Isotype Ctrl | PE | MOPC 21 | Biolegend | 400112 |
| Anti IgG1 κ Mouse, APC | APC | MOPC 21 | Biolegend | 400120 |

**Supplementary Table 5.**

| **Šídák's multiple comparisons test** | **Summary** | **Adjusted P Value** |
|---|---|---|
| Term BP vs. Term CAM | ns | 0.9993 |
| Term BP vs. First Trimester Decidua | ns | >0.9999 |
| Term BP vs. NT iPSC-dNK | ns | >0.9999 |
| Term BP vs. TGFβ iPSC-dNK | ns | 0.981 |
| Term BP vs. PBNK | ns | 0.1766 |
| Term BP vs. NK92 | ns | 0.3045 |
| Term CAM vs. First Trimester Decidua | ns | 0.5904 |
| Term CAM vs. NT iPSC-dNK | ns | 0.9813 |
| Term CAM vs. TGFβ iPSC-dNK | ns | 0.1422 |
| Term CAM vs. PBNK | ** | 0.0015 |
| Term CAM vs. NK92 | ** | 0.0095 |
| First Trimester Decidua vs. NT iPSC-dNK | ns | >0.9999 |
| First Trimester Decidua vs. TGFβ iPSC-dNK | ns | >0.9999 |
| First Trimester Decidua vs. PBNK | ns | 0.3269 |
| First Trimester Decidua vs. NK92 | ns | 0.5636 |
| NT iPSC-dNK vs. TGFβ iPSC-dNK | ns | 0.9389 |
| NT iPSC-dNK vs. PBNK | * | 0.05 |
| NT iPSC-dNK vs. NK92 | ns | 0.1618 |
| TGFβ iPSC-dNK vs. PBNK | ns | 0.942 |
| TGFβ iPSC-dNK vs. NK92 | ns | 0.9765 |
| PBNK vs. NK92 | ns | >0.9999 |
